# Comprehensive Metagenomic Profiling of Diverse Microbiomes

**DOI:** 10.1101/2025.02.13.638056

**Authors:** Balázs Kakuk, Ákos Dörmő, Ahmed Taifi, Tamás Járay, Gábor Kurucsai, Gábor Gulyás, István Prazsák, Zsolt Boldogkői, Dóra Tombácz

## Abstract

The canine gut microbiome serves as a key model for veterinary and human health research, but inconsistent findings arise due to methodological variations. This study presents a three-part dataset to clarify how DNA extraction, primer selection, and sequencing platforms influence microbial profiling. First, we performed ultra-deep sequencing of a single dog fecal sample using five DNA isolation kits, multiple library protocols, and four sequencing platforms (Illumina MiSeq/NovaSeq, ONT MinION, PacBio Sequel IIe), enabling direct comparisons of 16S rRNA and shotgun sequencing techniques. Second, we analyzed 40 fecal samples from eight co-housed dogs using Zymo High-Molecular-Weight (ZHMW) and Zymo MagBead (ZMB) extraction kits to assess longitudinal extraction-kit effects. Third, we evaluated three 16S primer systems (standard ONT, PacBio, and modified ONT with degenerate bases) using synthetic mock communities and human/canine fecal samples to quantify primer biases. By integrating synthetic and biological replicates, this dataset provides a standardized resource for benchmarking bioinformatics pipelines and improving cross-study comparability. The study generated 75.31GB of new sequencing data: 43.451GB from ZHMW-ZMB comparisons, 22.611GB for primer evaluations, and 9.191GB from the single-sample analysis. Combined with 31.51GB of prior data, the total dataset exceeds 1061GB, including all analytical outputs. These resources enhance methodological transparency and accuracy in canine gut microbiome research across diverse laboratory workflows.

## Context

Microbiome research plays an essential role in understanding the functions of microbial communities in health, disease, and environmental interactions. However, it faces persistent technical challenges arising from the lack of universal standards in DNA isolation, library preparation, sequencing, and bioinformatics workflows—all of which can introduce biases that affect reproducibility and comparability across studies [1]. DNA extraction is a particularly influential step; different kits vary in their efficiency at lysing microbial cells or retaining certain taxa, potentially altering observed community profiles [2,3]. Researchers must then choose between targeted 16S rRNA gene amplicon sequencing and whole-genome shotgun (WGS) sequencing, each of which balances cost, resolution, and coverage [4,5]. Sequencing platforms further complicate matters: while short-read sequencing (SRS) technologies (e.g., Illumina) are cost-effective and high-throughput, they struggle with highly repetitive or complex regions; conversely, long-read sequencing (LRS) platforms [Oxford Nanopore Technologies (ONT), Pacific Biosciences (PacBio)] enable more complete assemblies but require higher per-base costs and specialized protocols [6–9].

One of the most critical sources of bias in 16S rRNA gene studies is primer design. Conventional primers can fail to amplify essential taxa if mismatches occur in the conserved regions they target. Two recent studies underscore this issue: Matsuo et1al. (2021) demonstrated that the “27F” primer in the ONT 16S Barcoding Kit underrepresented *Bifidobacterium* and proposed degenerate primers to address this shortfall [10]. Waechter et1al. (2023) later validated these degenerate primers in a large human fecal dataset, reporting improved capture of microbial diversity and reduced taxonomic biases [11]. Together, these findings suggest that even minimal primer modifications can significantly enhance community profiling, particularly at the species level.

While considerable research focuses on human microbiomes, dogs (*Canis lupus familiaris*) serve as a valuable model with direct implications for veterinary and human health [12,13]. Early canine microbiome studies, predominantly using short-read sequencing, revealed how diet, host genetics, and environment influence gut communities [14]. More recently, long-read platforms have enabled near-complete bacterial genome assemblies from canine feces, providing deeper functional and taxonomic resolution [15]. Nevertheless, key questions remain about how best to integrate extraction kits, primer designs, and sequencing technologies to comprehensively characterize the canine gut microbiome.

With this dataset, we address these methodological challenges through three complementary analyses, each designed to clarify different aspects of workflow variability:

### Multi-platform, Single-Canine Fecal Evaluation

We analyzed the fecal sample from a single dog using multiple DNA extraction kits, library preparation methods (amplicon-and WGS-based), and sequencing platforms (SRS and LRS). To our knowledge, no prior study has subjected a single fecal sample—or any microbiome sample—to such a broad range of metagenomic methods: five DNA isolation kits, seven library preparation techniques, and four sequencing platforms. This unprecedented approach enables direct cross-comparison of short-and long-read data, as well as amplicon versus WGS strategies, within the same biological material.

### Forty Canine Fecal Samples (ZHMW vs. ZMB)

Next, we analyzed 40 dog fecal samples from a longitudinal and familial study. We compared two widely used DNA extraction kits—Zymo High-Molecular-Weight (ZHMW) and Zymo MagBead (ZMB)—chosen for their distinct mechanisms: ZHMW preserves longer DNA fragments (optimized for LRS), while ZMB employs a streamlined magnetic bead workflow favored for routine extractions. This setup highlights how extraction protocol choice impacts taxonomic composition at scale in a real-world canine cohort.

### Primer Comparison in Synthetic and Biological Samples

Finally, we evaluated three 16S primer sets—(A) standard ONT (V1–V9), (B) PacBio (PB) full-length 16S, and (C) modified ONT (MONT) with degenerate bases—applied to mock communities (Zymo D6300, D6331, D6323) and real fecal samples (human and canine). While degenerate primers have improved *Bifidobacterium* detection, the PB primer set has not been systematically compared to ONT or MONT protocols. This direct comparison quantifies primer-induced biases and identifies opportunities to refine microbiome profiling accuracy.

By integrating these investigations, our dataset provides a multifaceted perspective on how methodological decisions—DNA extraction, primer design, and sequencing platform—shape insights into the canine gut microbiome and broader microbial communities. This work builds on existing research addressing primer bias (e.g., *Bifidobacterium* underrepresentation) and aims to guide standardization and optimization of microbiome workflows for veterinary and translational studies.

## Data Description

In this Data Note, we present 106.761GB of raw metagenomic sequencing data along with their analytical outcomes. This total comprises 31.51GB from our previously published work [16], 9.191GB from a new Dog_M0 sample (sequenced using NovaSeq and MinION), 43.451GB generated from eight co-housed dogs (Serteperti cohort), and 22.611GB allocated to the primer comparison (including three synthetic and two biological samples). Table 1 summarizes the overall sequencing yields and other basic dataset information.

**Table 1.** Basic information about the datasets.

| Study phase | Source | Sample type | Dog No. | # of samples | # of replicates | DNA isolation method | Type of library | Platform | Gigabases (raw) | ENA Project |
| --- | --- | --- | --- | --- | --- | --- | --- | --- | --- | --- |
| Single-canine, Multilevel comparison | Our previous study [16] | Single dog, 2 samples | M0 | 1 | 4x4 | I, MN, Q, ZHMW | WGS | MiSeq | 13.72 | PRJEB59610 |
|  | Our previous study [16] |  | M0 | 1 | 4x4 | I, MN, Q, ZHMW | V1-V2 | MiSeq | 0.24 | PRJEB59610 |
|  | Our previous study [16] |  | M0 | 1 | 4x4 | I, MN, Q, ZHMW | V1-V3 | MiSeq | 0.96 | PRJEB59610 |
|  | Our previous study [16] |  | M0 | 1 | 4x4 | I, MN, Q, ZHMW | V3-V4 | MiSeq | 0.26 | PRJEB59610 |
|  | Our previous study [16] |  | M0 | 1 | 4x4 | I, MN, Q, ZHMW | V1-V9 | MinION | 15.90 | PRJEB59610 |
|  | Our previous study [16] |  | M0 | 1 | 4x4 | I, MN, Q, ZHMW | V1-V9 | Sequel | 0.43 | PRJEB59610 |
|  | This study |  | M0 | 2 | 1 | ZMB | WGS | NovaSeq | 5.41 | PRJEB75753 |
|  | This study |  | M0 | 2 | 1 | ZMB | WGS | MinION | 3.78 | PRJEB75753 |
| DNA isolation kit comparison | This study | Eight dogs | F1-F4 & M1-M4 | total of 40 | 3x3 | ZHMW, ZMB | V1-V9 | MinION | 43.45 | PRJEB82097 |
| Primer comparison | This study | Single dog, 1 sample | F5 | 1 | 3x3 | ZMB | V1-V9 | MinION | 22.61 | PRJEB85420 |
|  | This study | Single human sample | NA | 1 | 3x3 | ZMB | V1-V9 | MinION |  | PRJEB85714 |
|  | This study | MCS | NA | 1 | 3x3 | ZMB | V1-V9 | MinION |  | PRJEB85420 |
|  | This study | GMS | NA | 1 | 3x3 | ZMB | V1-V9 | MinION |  | PRJEB85420 |
|  | This study | FR | NA | 1 | 3x3 | ZMB | V1-V9 | MinION |  | PRJEB85420 |

### 1. Multi-platform, Ultra-Deep Sequencing of a Single Canine Feces Sample

#### Study Overview

Expanding upon our earlier study [16], we performed ultra-deep, multi-platform sequencing on a single fecal sample from a male Pumi dog (Dog_M0, nicknamed “Toti”; see Supplementary Table S1 for metadata), to evaluate how DNA extraction kits and library preparation methods influence microbial profiling across short-and long-read sequencing platforms. Supplementary Table S2 shows the read counts and other information on the previously published Illumina 16S and ONT 16S V-region sequencing data. In addition to the sample analyzed in our prior work, this new dataset incorporates additional WGS runs—including Illumina NovaSeq and ONT MinION—enabled by a fifth DNA isolation protocol (Figure 1). Supplementary Table S3 presents the sequencing yields for the two novel datasets.

**Figure 1.**
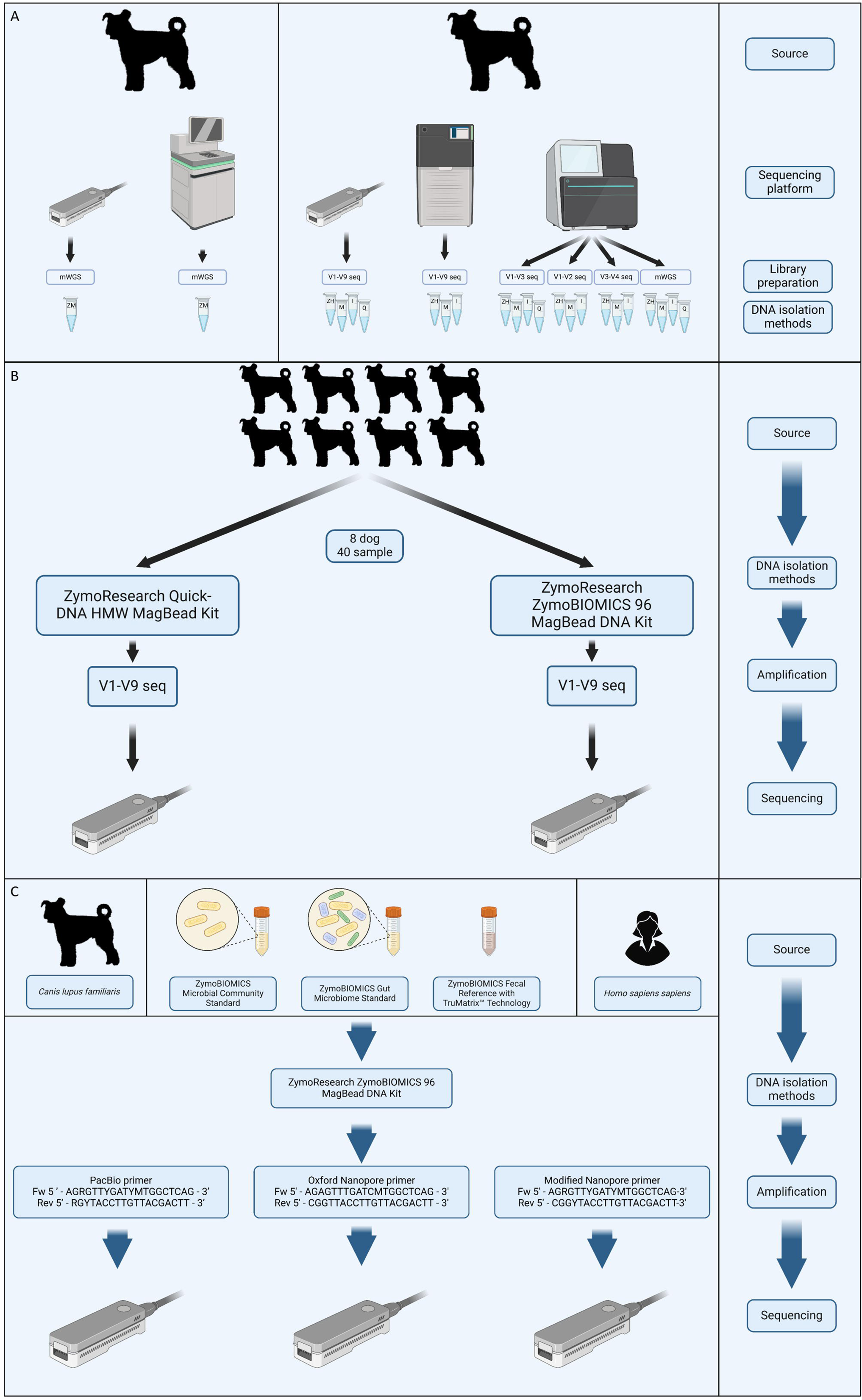
Experimental arrangement.

#### Experimental Arrangement

##### DNA Isolation Methods (4 previously used + 1 additional)

1. Qiagen QIAamp Fast DNA Stool Mini Kit (Q)
2. Invitrogen PureLink™ Microbiome DNA Purification Kit (I)
3. Macherey-Nagel NucleoSpin® DNA Stool Mini Kit (M)
4. Zymo Research Quick-DNA™ HMW MagBead Kit (ZHMW)
5. Zymo Research MagBead DNA Isolation Kit (ZMB) – newly introduced for additional WGS runs

The first four kits follow the protocol described by Gulyás et1al. [16]. For the new NovaSeq and ONT WGS data, we utilized the Zymo MagBead protocol. Each method differs in its lysis efficiency and capacity to preserve high molecular weight (HMW) DNA, which could influence taxonomic profiles.

##### Library Preparation Kits (16S and WGS)

1. Illumina DNA Prep (Document #1000000025416 v09)
2. ONT Rapid Sequencing 16S Barcoding Kit (SQK-RAB204)
3. ONT Ligation Sequencing Amplicons – Native Barcoding Kit 96 V14 (SQK-LSK114.96)
4. PacBio Full-Length 16S Library Prep (SMRTbell Express Template Prep Kit 2.0)
5. PerkinElmer NEXTFLEX® 16S V1–V3 Amplicon Seq Kit (Cat. #: 4202-03)
6. Zymo Research Quick-16S™ NGS Library Prep Kit (Ca.#: D6400)
7. IDT xGen™ DNA Lib Prep Kit (Cat. #: 10009822)
8. ONT Ligation Sequencing Kit V14 (SQK-LSK114)

The first four kits follow the protocol described by Gulyás et1al. [16]. For the new NovaSeq and ONT WGS data, we utilized the xGen™ DNA Lib Prep Kit and the ONT Ligation Sequencing protocol, respectively.

##### Sequencing Platforms

- Illumina MiSeq: Generated metagenomic data in prior runs as detailed in Gulyás et1al. [16].
- llumina NovaSeq 6000: Applied for ultra-deep WGS.
- Oxford Nanopore Technologies MinION: Used for V1-V9 16S rRNA and WGS runs.
- Pacific Biosciences Sequel IIe: Employed for full-length 16S (V1–V9) amplicon sequencing.

##### Data Processing

- 16S rRNA (V-region): Processed using EMU to generate taxonomic abundance profiles from amplicon kits targeting V1–V2, V1–V3, V3–V4 (Illumina), and V1–V9 (ONT/PacBio).
- WGS: Reads were classified with Kraken2 and sourmash for short-read (MiSeq, NovaSeq) and long-read (ONT) data.

By utilizing a diverse array of extraction kits, library preparation methods, and sequencing platforms on these two fecal samples, we establish a comprehensive framework to evaluate methodological biases and directly compare amplicon-and WGS-based strategies in canine gut microbiome research.

### 2. Throughout Comparison of Zymo MagBead and Zymo HMW Kits Using 40 Canine Feces Sample

#### Study Overview

This dataset comprises fecal samples collected over a one-year period from eight co-housed, purebred adult Pumi dogs at the Hungarian *Serteperti Pumi Kennel*, aimed at investigating temporal shifts and familial patterns in the gut microbiome. For each sample, DNA was isolated using both the Zymo High-Molecular-Weight (ZHMW) and Zymo MagBead (ZMB) kits, enabling direct comparison of these two extraction method. Supplementary Table S1 provides the details about the dogs, while Supplementary Table S4 contains information on the sequencing yields.

#### Experimental Arrangement

Cohort: Eight adult dogs (F1–F4, M1–M4), including five closely related individuals (father, daughter, granddaughter, and littermates), were housed in a shared environment at the Serteperti Pumi Kennel. This setup minimized variability in genetics, diet, and housing conditions. All dogs received the same diet (see Methods). Figure 2 shows the familial relationships of the five genetically related dogs in the kennel that were sampled in this project.

**Figure 2.**
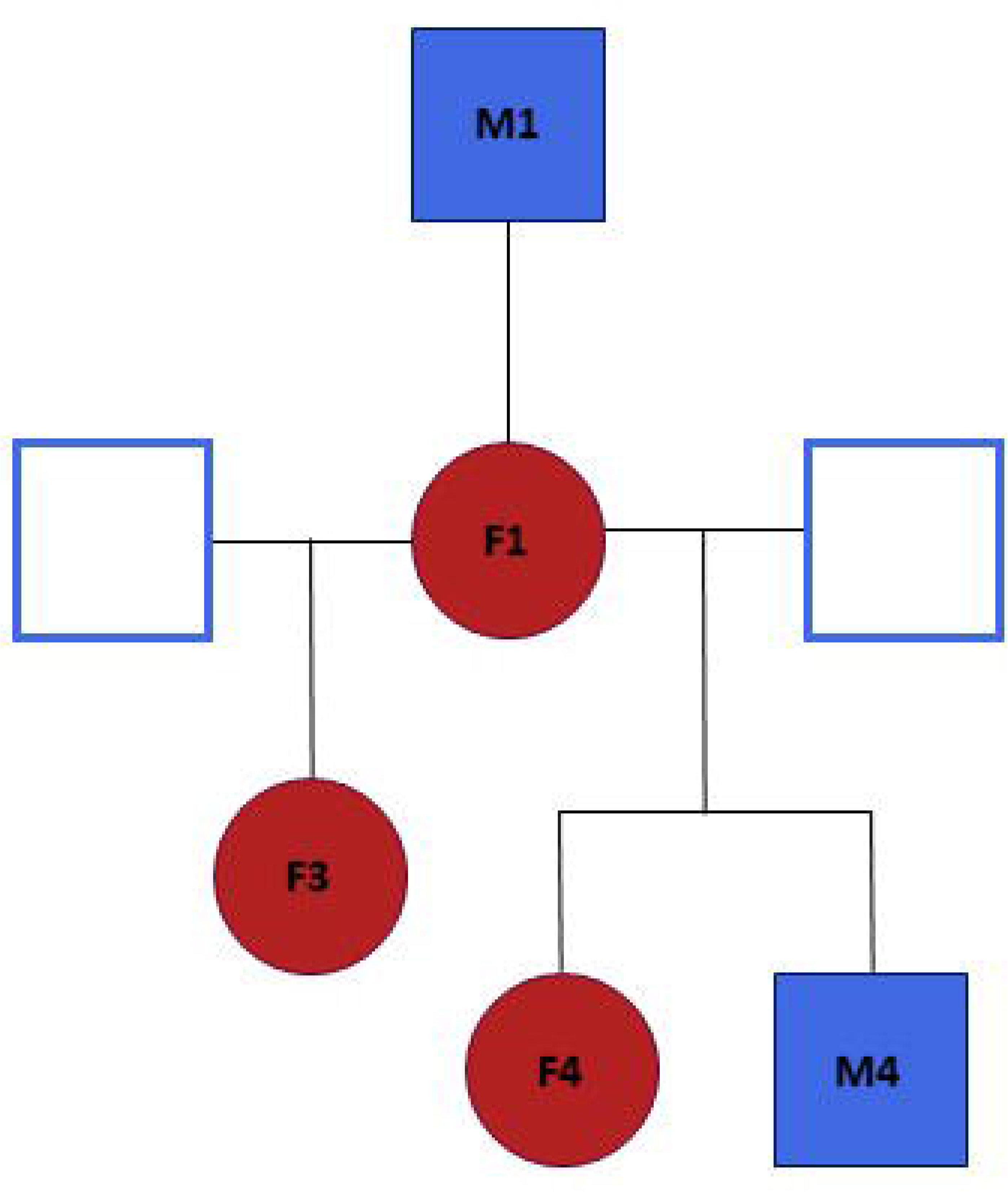
Familiar relationships among the dogs from the Serteperti cohort.

**Figure 3.**
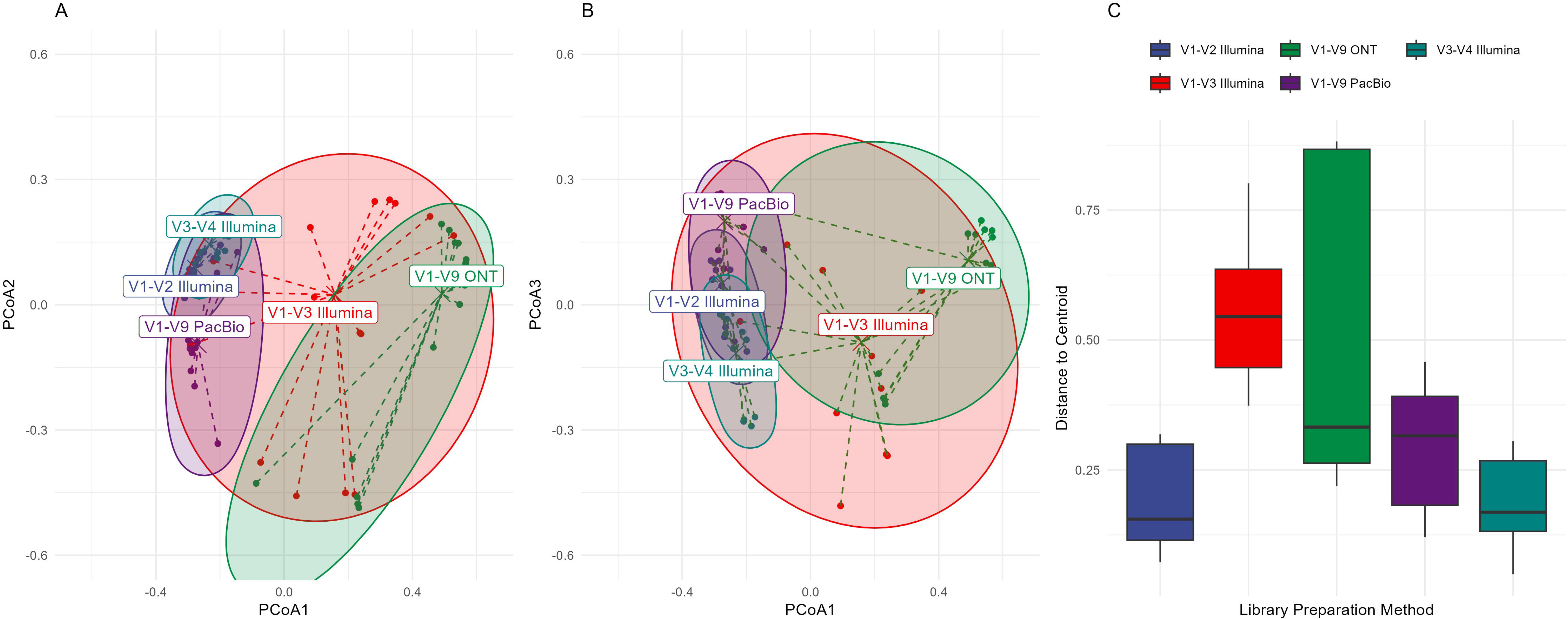
Group dispersions for different library preparation protocols in the first sampling of the single canine dataset. A PERMDISP2 analysis was employed to measure beta diversity differences among library methods. (**A**) Principal coordinates analysis (PCoA) of PCoA1 vs. PCoA2, where dashed lines connect each point to the centroid of its respective protocol; (**B**) PCoA1 vs. PCoA3, illustrating a similar dispersion pattern; and (C) Boxplots of distance-to-centroid for each library group. Ninety-five percent confidence ellipses in (A) and (B) depict group spread, while panel1(**C**) shows how tightly (or loosely) the samples cluster around their centroids.

**Figure 4.**
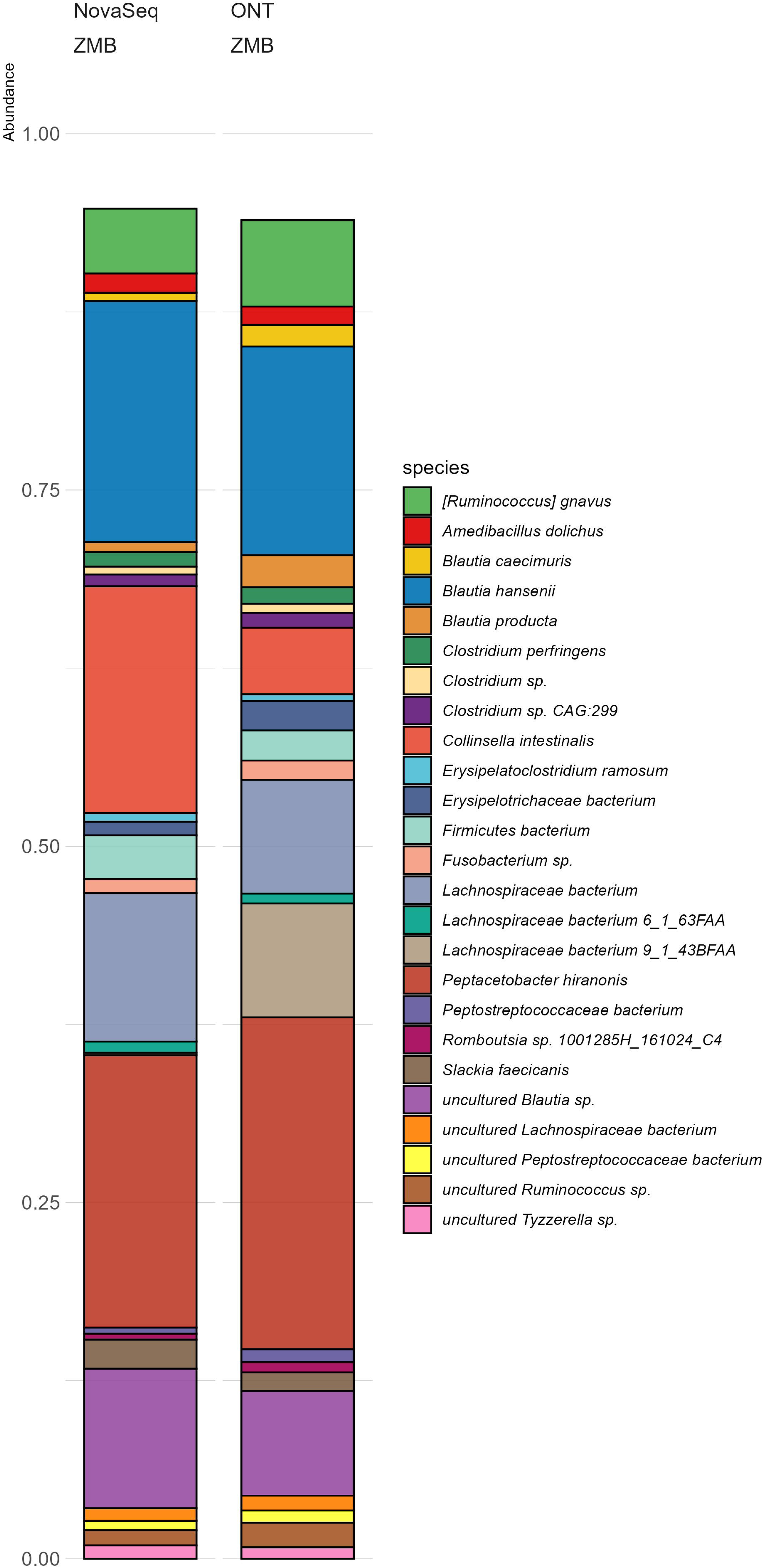
Prokaryotic species distribution in the second sampling of the single canine dataset based on NovaSeq and ONT whole-genome sequencing. Stacked bar plots show the top 25 prokaryotic species abundances. Unclassified reads were excluded from the relative abundance calculation. The Y-axis shows the proportional abundance for each species. Unfilled portion in each bar represents taxa outside the top125.

**Figure 5.**
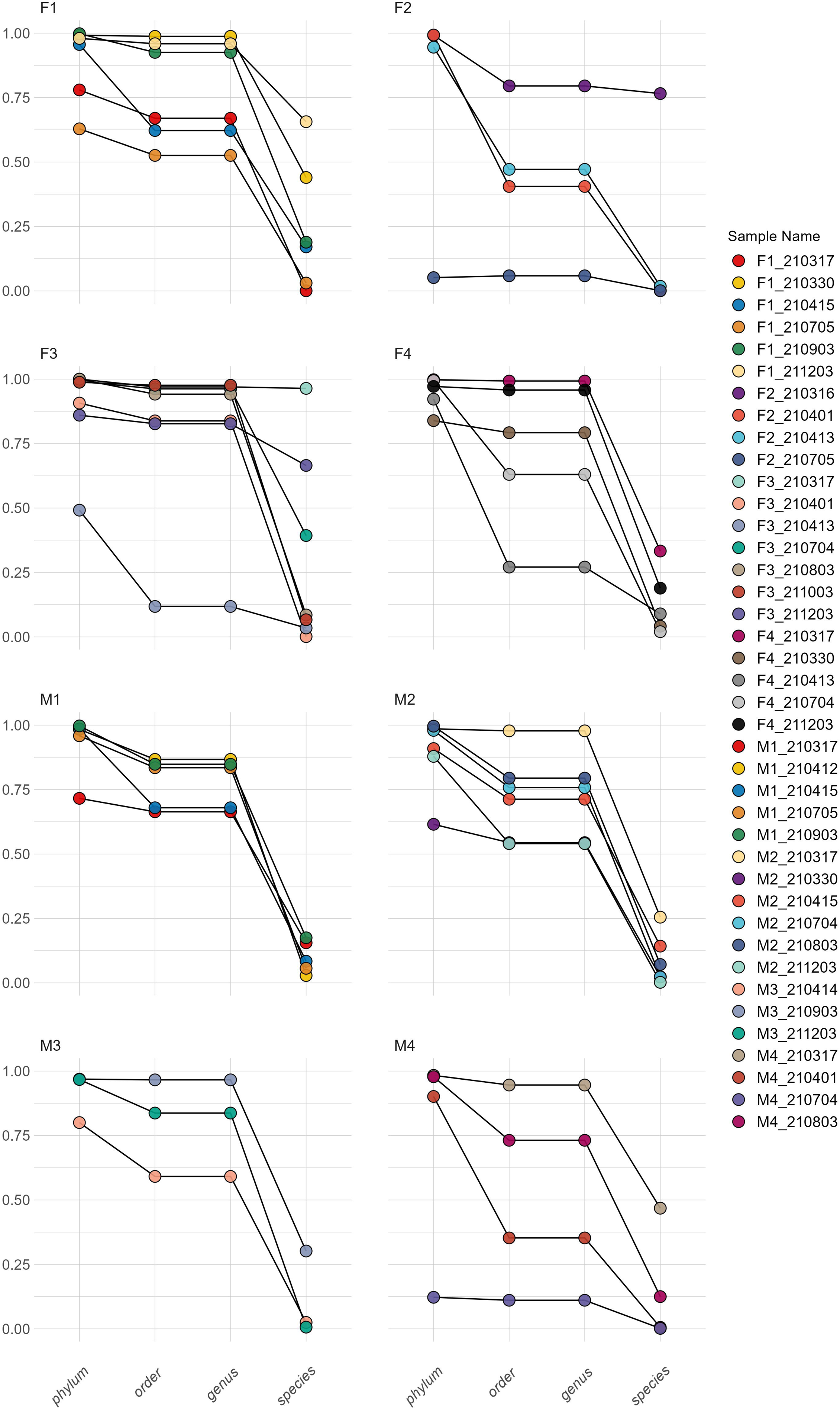
Taxon-abundance correlations between two DNA isolation methods for each sample across four taxonomic ranks. Line plots illustrating the correlation in terms of Pearson’s R˄2 values of the taxonomic abundance for each taxon between the ZMB and ZHMW kits. Each line, color-coded by sample name, connects a single sample’s *R*˄2 across taxon levels (phylum, order, genus, species), while panels (facets) separate samples by their source (dog).

**Figure 6:**
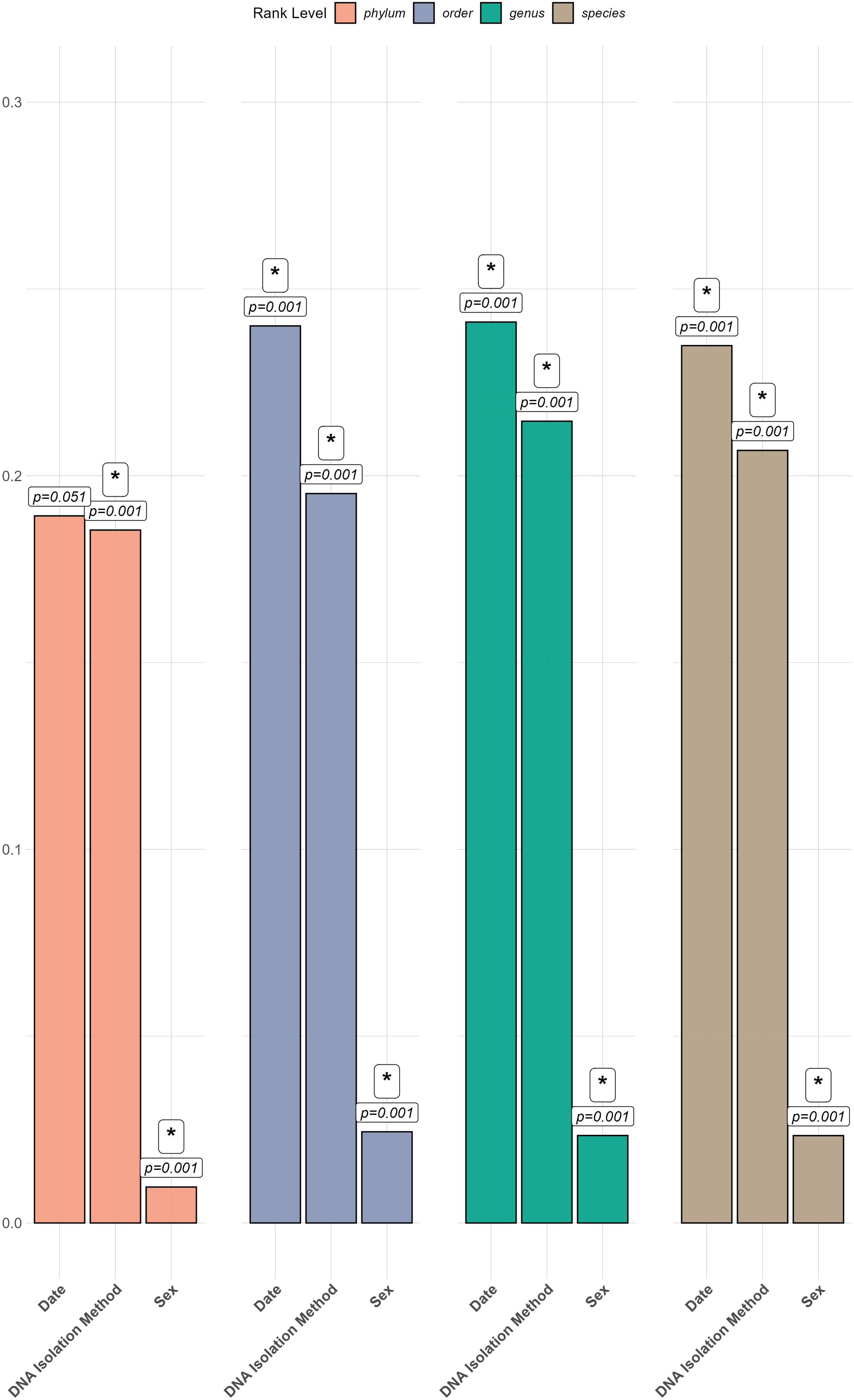
Contribution of DNA isolation kit, host (dog) sex, and collection date to total microbiome variability (species-level abundances) in the DNA method comparison on forty canine samples dataset. The y-axis on the bar plot of shows PERMANOVA *R*˄2 values at four taxonomic ranks (phylum, order, genus, species), illustrating how “DNA Isolation Method,” “Sex,” and “Date” each contribute to variability in the microbiome. Each facet corresponds to a different rank level, and bars within each facet represent the *R*˄2 value for one explanatory factor. Labeled p-values and significance indicators appear above the bars, highlighting whether each term is statistically significant at the respective rank.

**Figure 7.**
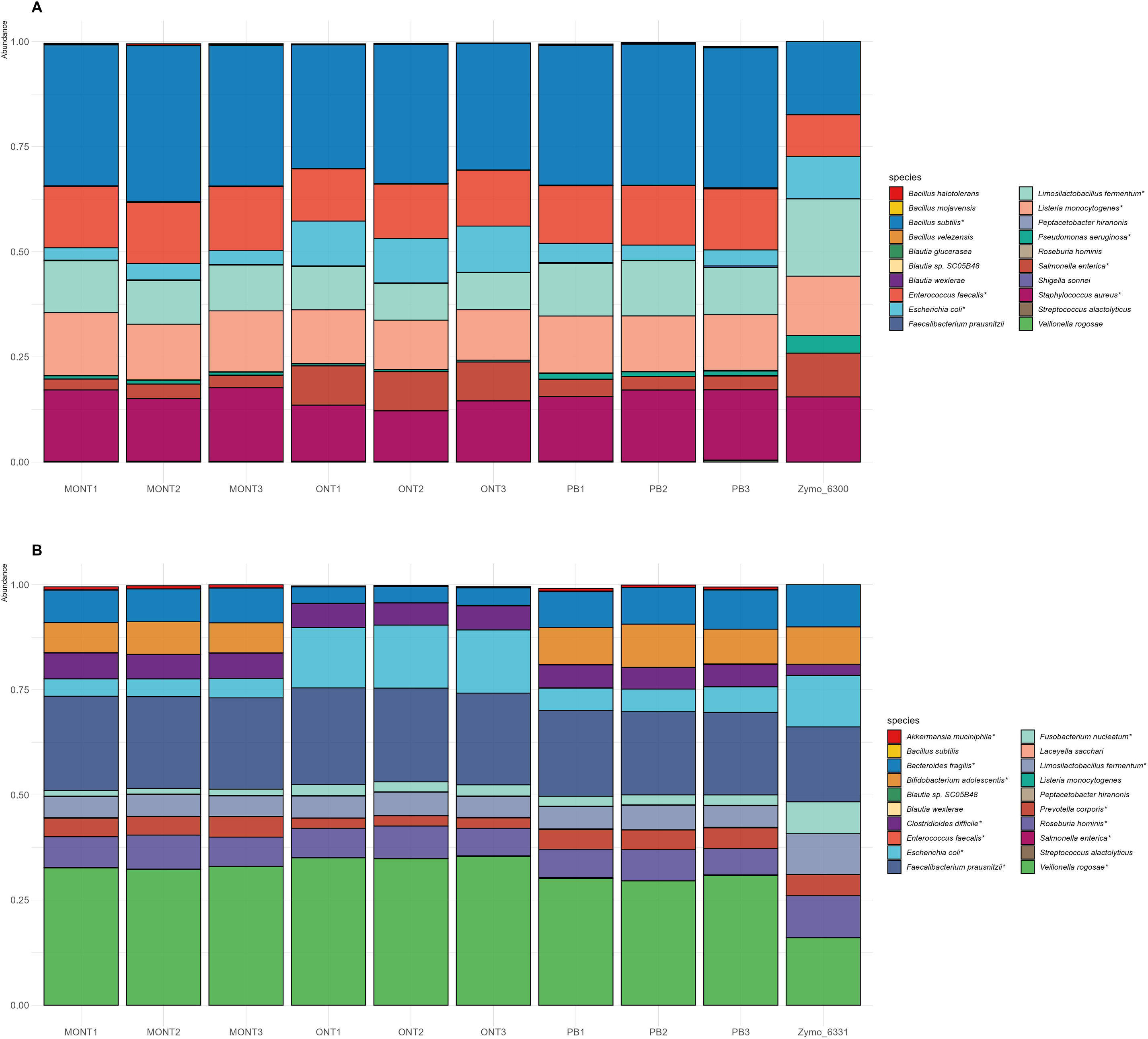
Theoretical vs. experimentally observed taxonomic composition in two Zymo Mock Community Standards using three 16S primer sets. Comparison of theoretical vs. experimentally observed taxonomic composition for two Zymo Mock Community Standards assessed by the three 16S primer sets (ONT – original Oxford Nanopore Technologies V1-V9 primer, MONT – Modified ONT primer and PB – PacBio primer). Panel **A)** shows the results for D6300 and **B**) for D6331. Each bar represents a species from the EMU-based classifications of the V1–V9 ONT sequencing, while the final bars on the right of each panel show the manufacturer’s stated (theoretical) composition. The Y-axis shows the proportional abundance for each species.

**Figure 8.**
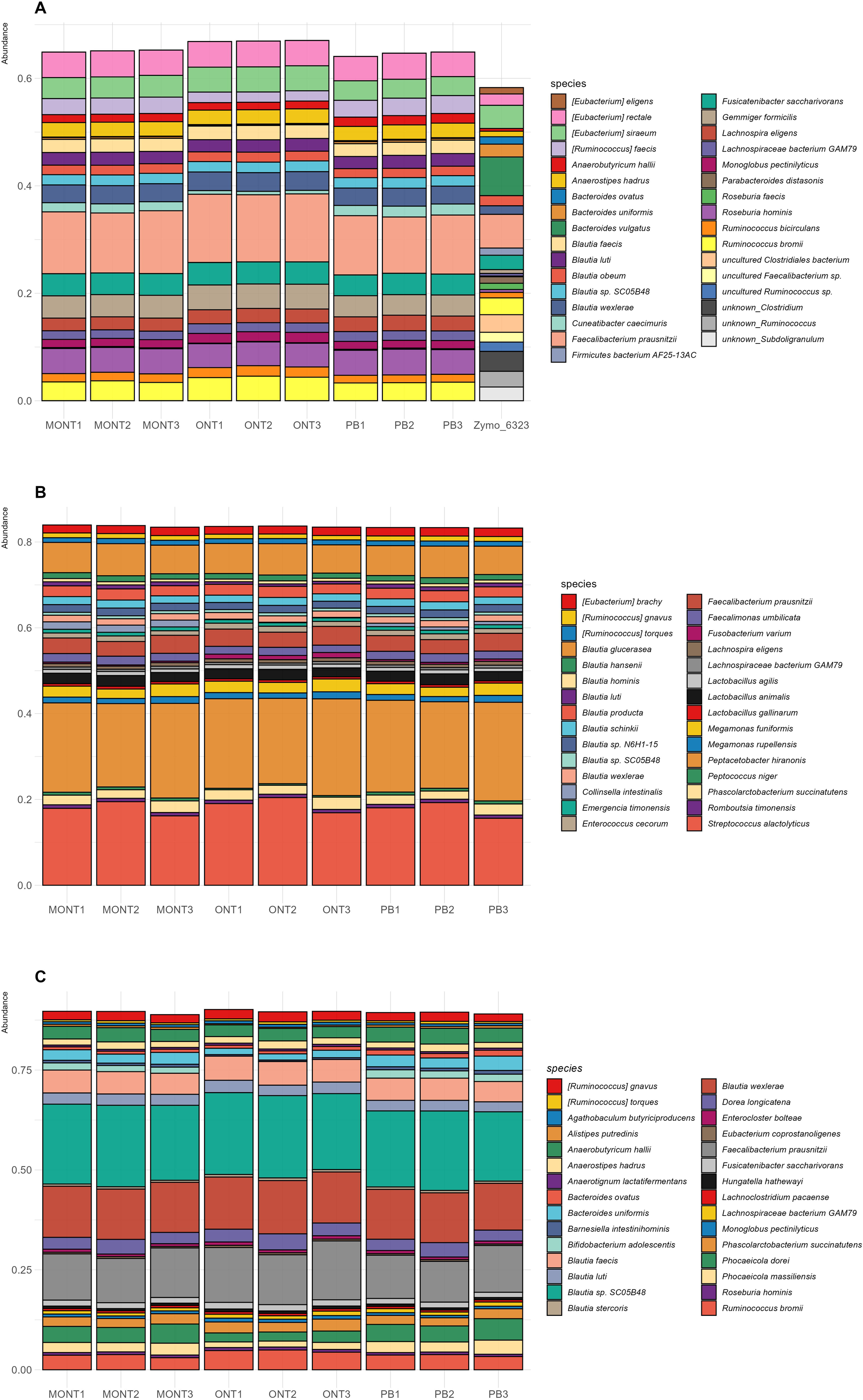
Comparison of the top130 taxa in Zymo FR D6323, canine (*C. lupus familiaris*) feces, and human feces under three 16S primer sets. (A) Zymo FR D6323 (with a WGS1+1Kraken12 reference composition shown), (B) *C. lupus familiaris* fecal sample, and (C) human fecal sample. All data were derived via EMUIZlbased classification of V1–V9 ONT sequencing, with each bar illustrating a species in the ampliconIZlderived profiles. **T**he three 16S primer sets used in the library preparation were ONT – original Oxford Nanopore Technologies V1-V9 primer, MONT – Modified ONT primer and PB – PacBio primer. The Y-axis shows the proportional abundance for each species. Unclassified reads were omitted from the relative abundance calculation. Unfilled portion in each bar represents taxa outside the top130.

Longitudinal Sampling: Fecal samples were collected at multiple time points over a 12-month period to capture routine microbiome shifts and gestational events (Supplementary Table S1).

Dual Extractions: For the 40 fecal samples, fecal material was divided for parallel DNA extraction using the Zymo High-Molecular-Weight (ZHMW) and Zymo MagBead (ZMB) kits. Resulting DNA extracts were sequenced on the Oxford Nanopore MinION platform, targeting the full-length 16S rRNA gene (V1–V9 region).

Analysis: Sequencing data were basecalled using Dorado, followed by taxonomic profiling with EMU. A comprehensive R workflow was developed for downstream analysis and visualization. Supplementary Table S2 provides metadata for the 40 matched-extraction samples, including:

- Dog ID: F1–F4 (female), M1–M4 (male)
- Sex
- Collection Date (monthly intervals to reflect temporal variation)
- Extraction Method (ZHMW or ZMB)

### 3. Comparison of Three Different 16S Primer Sets in Synthetic and Biological Samples

#### Study Overview

In this section of the study, we evaluated the performance of three distinct primer systems across five sample types:

- The standard ONT 16S V1–V9 protocol,
- A PacBio (PB) full-length 16S rRNA approach, and
- A modified ONT (MONT) protocol incorporating degenerate bases.

While previous studies have demonstrated the efficacy of degenerate primers in improving *Bifidobacterium* detection with ONT [10], the PB primer system has not yet been directly compared to ONT or MONT. This comparison addresses a critical gap in understanding 16S primer biases. Supplementary Table S5 contain information about the sequencing yields in this dataset.

#### Experimental Arrangement

##### Primer Sets

- ONT: Targets the V1–V9 region of the 16S rRNA gene.
- PB: Full-length 16S rRNA gene (V1–V9).
- MONT: Incorporates degenerate bases into the ONT protocol.

##### Sample Sources

- Zymo D6300: A simplified synthetic mock community containing eight bacterial strains.
- Zymo D6331: A complex synthetic mock community with 18 bacterial species.
- Zymo D6323: Standardized human fecal reference (FR).
- Canine fecal sample: Collected from a female Pumi dog “F5” (see metadata in Supplementary Table S1).
- Human fecal sample: Anonymous female donor (aged 53, from Hungary, collected in Székesfehérvár).

Each of the five sample types was combined with the three primer sets and sequenced on the ONT MinION platform in triplicate, yielding 45 total sequencing runs. Basecalling was performed using Dorado, and taxonomic assignments were generated from the FASTQ files using the EMU pipeline. Final abundance profiles were processed in R with custom in-house scripts (see Materials and Methods for details).

## Analyses

### 1. Hybrid, Ultra-Deep Sequencing of Feces from the Same Canine Source

#### 1.A. Comparative Taxonomic Profiles Using Multiple V-Regions and Platforms

Expanding upon our earlier findings [16], we generated additional Illumina WGS and Oxford Nanopore (ONT) WGS data from a new fecal sample from the same donor (Dog_M0). Figure 3 illustrates how library preparation methods and sequencing platforms (including Illumina NovaSeq and ONT WGS) influenced species-level taxonomic abundances. Notably, *Peptacetobacter hiranonis* remained highly abundant in both ONT (∼23%) and NovaSeq (∼19%), whereas *Collinsella intestinalis* exhibited a marked increase in NovaSeq (15.9% vs. 4.7% in ONT). Other taxa, such as *Blautia producta*, declined from 2.2% (ONT) to 0.7% (NovaSeq), highlighting variable performance of short-and long-read WGS for individual species. Minor taxa also showed platform-specific absences or overrepresentations, underscoring the need for cross-method validation.

Additionally, we assessed the impact of DNA isolation method and library preparation on community variance using PERMANOVA (Supplementary Figure S2). Inclusion of the Qiagen (Q) kit—previously associated with lower-quality DNA—introduced substantial statistical noise, with Q samples frequently appearing as outliers (Supplementary Figure S2A). Consequently, DNA isolation and library preparation each explained ∼38% of the total variance (R² ≈ 0.385 and 0.382, respectively). However, after excluding Q samples (Supplementary Figure S2B), these factors accounted for larger variance shares (R² ≈ 0.449 for isolation method, R² ≈ 0.412 for library prep), demonstrating that suboptimal DNA extractions can obscure true method-driven differences.

#### 1.B. Illumina NovaSeq vs. ONT WGS Using Zymo MagBead

Expanding on part (A), Figure 3 highlights the top 25 species abundances in the Zymo MagBead extracts for both ONT and NovaSeq. Taxa like *Fusobacterium* or *Coprococcus* display notable differences between platforms, with some species entirely absent in one dataset. These results confirm that platform-specific biases can lead to divergent profiles and highlight the importance of cross-platform comparisons in single-sample metagenomic studies

#### 1.C. Viral Community Analyses

While bacterial diversity was our primary focus, we also profiled viral taxa from the whole-genome shotgun (WGS) datasets to compare the five DNA isolation kits. Supplementary Figure S3 shows the top 30 viral species detected. Although viral taxa constituted a minor fraction overall, we observed notable discrepancies in the viral communities across multiple metagenomic WGS (mWGS) runs from two fecal samples of the same dog (“Dog_M0”), reflecting methodological differences and possibly temporal variation. For example, viruses such as *Kingevirus communis* and *Lahndsivirus rarus* dominated newer NovaSeq and ONT datasets but were minimally present or absent in earlier MiSeq runs. Conversely, viruses like *Erskinevirus EaH2, Natevirus nate*, and *Microbacterium phage Pumpernickel* exhibited higher relative abundances in older MiSeq data. These shifts may arise from temporal changes in the dog’s gut virome, combined with platform-and library prep-specific biases (e.g., sequencing depth, detection sensitivity, or bioinformatic classification pipelines).

### 2. Comparison of Zymo MagBead and Zymo HMW Kits Using 40 Canine Fecal Samples

#### 2.A. Assessment of Contribution of Factors to the Variability in Taxon Abundances

We next compared Zymo High-Molecular-Weight (ZHMW) and Zymo MagBead (ZMB) DNA extraction methods using 40 canine fecal samples from eight co-housed dogs, with each dog sampled longitudinally. After blocking for dog identity and adjusting for sex and collection date as covariates in PERMANOVA, DNA isolation method emerged as a primary driver of variability across all taxonomic levels. At the phylum level, extraction method was highly significant (p1=10.001), explaining ∼18.5% of the variance, while sex was also significant (p1=10.001) but contributed only ∼1%. Collection date showed borderline significance (p1≈10.051). At finer taxonomic resolutions (order, genus, species), DNA isolation method remained the dominant factor (p1=10.001; ∼20–21% variance explained), while sex and date became more influential (p1<10.01), contributing ∼2–2.4% and ∼19–24% variance, respectively. These findings highlight that extraction kit choice strongly impacts observed gut microbiome composition, particularly at higher taxonomic ranks, where subtle methodological biases may amplify differences between ZHMW and ZMB workflows.

#### 2.B. Correlation Analyses of Taxon Abundances

To further evaluate consistency between ZHMW and ZMB extractions, we calculated Pearson correlations (R²) for the top 20 taxa at each taxonomic rank per sample. Figure 5 summarizes these correlations. On the Phylum level most samples showed high correlation (R² > 0.9), indicating broad-scale compositional agreement. However, outliers (e.g., F2-210705) exhibited very low R² (∼0.05), suggesting kit-dependent anomalies. On lower taxonomic levels (Order, Genus and Species) correlation ranges broadened, with some samples retaining high alignment and others dropping to near-zero values, particularly at the species level. For example, M3-211203 displayed R²1≈10.0003, reflecting pronounced extraction-driven biases for specific fine-level taxa.

Overall, these correlation analyses corroborate the PERMANOVA results, demonstrating that DNA isolation method strongly influences community profiles. While phylum-level distributions often align closely, finer taxonomic ranks diverge depending on sample-specific factors.

### 3. Comparison of Three Different V1-V9 Primer Sets in Synthetic and Biological Samples

#### 3.A. Overview and Rationale

Finally, we investigated three 16S rRNA primer systems—standard ONT (V1–V9), PacBio full-length 16S (PB), and a modified ONT (MONT) with degenerate bases—to address biases such as *Bifidobacterium*underrepresentation. We tested each primer set on five sample types (two synthetic mocks, a standardized fecal reference, a canine fecal sample, and a human fecal sample) to gauge primer-induced variation across controlled and real-world contexts.

#### 3.B. Zymo D6300 and D6331 (Mock Communities)

In the simpler D6300 mock, certain species (e.g., *Bacillus subtilis*) were consistently overrepresented, while others (e.g., *Limosilactobacillus fermentum*) appeared underrepresented. Some species that were expected to be absent appeared in low abundances, which could be due to classifier misassignments or minor contamination. The more complex D6331 standard displayed pronounced differences in key taxa such as *Bifidobacterium adolescentis*, detected at near-expected abundance (8–10%) with MONT or PB, but nearly zero with standard ONT. Conversely, *Escherichia coli* often showed higher in the ONT data and lower in PB or MONT, underscoring that each primer set selectively favors or suppresses particular taxa.

#### 3.C. Fecal Reference (FR), Human, and Canine Samples

When comparing the Zymo fecal reference (FR) to a real human sample and a canine sample under V1–V9 16S sequencing, we observed substantial primer-driven variations. Several *Bacteroides* species expected in FR were underrepresented or missing in the amplicon data, while *Eubacterium rectale, Faecalibacterium prausnitzii*, and *Roseburia hominis* exceeded reference proportions. It is important to emphasize, however, that the reference composition was generated via the manufacturer’s WGS1+1Kraken2 pipeline. Consequently, these comparisons reflect differences among multiple workflows rather than evaluating the 16S amplicon data against a universally accepted “gold standard.” In some instances (e.g., *Bifidobacterium, Ruminococcus*), FR’s relative abundance aligned more with the dog than the human sample. This finding underscores that primer selection and sequencing platform factors can yield unexpected overlaps in taxonomic composition, necessitating caution when interpreting “reference” fecal samples as definitive benchmarks.

## Discussion

### Comparing ONT and Illumina (NovaSeq) WGS

Our results indicate that sequencing platform and library preparation can generate substantial discrepancies in species-level abundances—even when applied to the same fecal sample. For instance, *Collinsella intestinalis* showed significantly higher representation in NovaSeq data than in ONT WGS, whereas *Blautia producta* was comparatively depleted in NovaSeq. These findings align with previous observations that short-read and long-read technologies differ in their capacity to detect or accurately classify certain taxa [17]. While both platforms captured the dominant community members consistently (e.g., *Peptacetobacter hiranonis*), minor taxa frequently shifted or even disappeared, emphasizing the need for cross-platform validation. Researchers aiming for comprehensive community resolution may therefore benefit from combining short-and long-read datasets or at least acknowledging where one technology’s biases may overshadow subtle population differences.

### Impact of Zymo MagBead vs. Zymo HMW Kits on Canine Fecal Profiles

In the eight dog, forty-sample canine cohort, DNA isolation method emerged as the most significant factor at multiple taxonomic ranks, surpassing even individual dog identity, sex, and collection date in explaining community variance. While sex and date became relevant at finer resolutions (e.g., genus/species-level analyses), these biological covariates contributed comparatively small fractions of the variance. The consistent effect of extraction protocol (MagBead vs. HMW) underscores that kit selection can dramatically influence observed microbial distributions—potentially masking or exaggerating shifts in low-abundance taxa. Our correlation analyses further revealed that high-level taxonomic assignments (e.g., phylum) remain relatively consistent between kits, but genus-and species-level detections diverged significantly for certain samples. This outcome highlights the importance of matching DNA extraction protocols to study objectives, particularly when investigating fine-scale community structures or identifying minor, extraction-sensitive constituents.

### Primer Comparisons Across Synthetic and Host-Associated Samples

By evaluating three 16S primer sets on synthetic microbial standards and real fecal samples (human and canine), we revealed primer-induced biases that are often overlooked in studies relying on a single protocol. Modified ONT (MONT) and PacBio (PB) primers demonstrated superior accuracy in reconstructing mock community compositions, particularly for *Bifidobacterium*, which standard ONT primers systematically underrepresented. This enhanced detection aligns with recent findings that minor adjustments to primer sequences - such as degenerate bases or rearrangements - can significantly improve coverage of key gut taxa. Critically, these improvements persisted in real fecal samples, where MONT and PB primers either matched or exceeded reference abundances for genera prone to underamplification. However, each primer set exhibited distinct trade-offs, highlighting the need to validate 16S workflows using both mock communities (to benchmark “ground truth” compositions) and host-associated samples (where strain diversity, host diet, or other confounders may obscure primer biases).

### Concluding Remarks

Taken together, these analyses demonstrate how every step in a microbiome study - from DNA extraction to primer selection, and from sequencing platform choice to data processing - can yield noticeably different community profiles. Although broad phylum-level patterns often remain relatively stable, species-level composition may fluctuate dramatically, revealing biases that can skew interpretations of host-microbe interactions. Researchers may mitigate these effects by (i) selecting extraction kits validated for their sample types, (ii) testing primer sets against mock communities containing relevant target taxa (e.g., *Bifidobacterium*), and (iii) considering cross-platform comparisons or hybrid assemblies for high-resolution investigations. Ultimately, our study underscores the importance of rigorous method selection, thorough reporting of protocols, and ongoing benchmarking initiatives to ensure reproducible and accurate assessments of the canine gut microbiome—and potentially beyond, to other host or environmental systems.

## Reuse Potential

### Single-Canine, Multi-Platform Dataset

The multi-platform, multi-extraction dataset derived from a single fecal sample provides a valuable standard for comparing amplicon– versus shotgun-based approaches, short– versus long-read sequencing, and diverse library preparation protocols. Researchers can leverage these data to quantify method-induced biases, validate new classification pipelines against thoroughly cross-checked ground-truth data, and explore how each step—from DNA isolation kit to sequencing platform—alters perceived community composition. Because this resource includes both variable (V) region and WGS sequencing data, it is especially valuable for laboratories aiming to optimize workflows across different project types (e.g., rapid 16S screens, deep WGS-, or even virome studies). By serving as a foundational reference, the dataset accelerates method standardization and supports robust cross-platform comparisons in canine microbiome research and beyond.

### DNA Method Comparison on Forty Canine Samples

This subset of 40 fecal samples, each extracted in parallel using ZHMW and ZMB protocols, serves as a valuable reference for evaluating how extraction kit choice influences microbiome profiling in canine feces. The cohort includes eight Pumi dogs, five of which are direct relatives, with well-documented diet and housing conditions. This unique dataset allows for multiple comparative analyses:

1. Longitudinal investigations, as each dog was sampled at least three times, with most having 5–8 samples collected over time;
2. Cross-study comparisons, enabling integration of these data with other metagenomic studies, particularly those using long-read sequencing or dietary interventions;
3. Aging-microbiome research, facilitated by the inclusion of additional samples from the study, covering a total of 10 dogs with 43 samples spanning an age range of 2.1 to 14.8 years. In addition, researchers can evaluate their pipelines against technical replicates, identifying method-induced biases at various taxonomic levels. The controlled environmental variables (shared diet and housing) and paired extractions per sample make this an ideal testbed for refining DNA isolation strategies and quantifying how kit changes affect apparent microbial diversity over time.

### V1-V9 Primer Comparison on Synthetic and Biological Samples

The dataset contrasting standard ONT, PacBio, and modified ONT (MONT) primer sets serves as a key resource for examining primer-induced biases, particularly for underrepresented taxa like *Bifidobacterium*. By including synthetic microbial mixtures (Zymo D6300, D6331, D6323) with known compositions, researchers can directly calculate detection statistics - such as true positives, false positives, and correlations with theoretical abundances - and evaluate how each primer set amplifies or misrepresents specific taxa. Integrating these mock community data with real fecal samples (canine and human) enables a nuanced view of how primer choice and community complexity jointly affect taxonomic profiling. The dataset provides a robust platform for testing and validating bioinformatics pipelines, including primer trimming, taxonomic assignment, and error-correction routines under both controlled and natural conditions. Ultimately, by bridging known and unknown microbial compositions, the dataset empowers researchers to refine 16S rRNA protocols for more robust, bias-aware profiling across diverse sample types.

Overall, this comprehensive data source supports comparisons of microbial profiles across sample types, DNA isolation kits, sequencing library preparations, primers, platforms, and analytical methods.

## Materials and Methods

The detailed workflow of the study is illustrated in Figure 1. Both the DNA isolation and library preparation were conducted strictly according to the protocols provided with the kits. Detailed descriptions of these procedures are outlined below.

### Sample Sources

The details about the dogs used in the study can be found in Supplementary Table S1.

#### 1. Single Canine Fecal Sample

A single fecal sample was collected from an adult *Canis lupus familiaris* (Dog_M0). The dog was privately owned, and the feces were obtained shortly after defecation to minimize environmental contamination. The collected material was immediately stored at −201°C, then placed to −801°C within a day. The sample (1.01g) was subdivided for multiple DNA extraction and library preparation methods (metagenomic whole-genome sequencing as well as 16S rRNA amplicon sequencing). The sample was divided into portions according to the requirements of the different extraction kits. To minimize bias, the portions were randomly assigned rather than being allocated based on their position within the original sample. Due to the limited sample size, homogenization was not performed. This approach aligns with the findings of Liang et al. [18], who demonstrated that while homogenization is essential for metabolomic analyses, it is not necessary for microbiome studies. Supplementary Table S2 contains the sequencing read counts and barcodes for the previously published data [16] and Supplementary Table S3 contains information about the newly sequenced datasets from this dog.

#### 2. Forty Dog Fecal Samples

Fecal samples were collected from eight co-housed adult dogs from the Serteperti Pumi Kennel (Dány, Hungary) over a one-year period (the Serteperti cohort). Each dog produced multiple time-point samples, and 40 of these were selected for a direct comparison of two DNA isolation methods (Zymo High-Molecular-Weight vs. MagBead). Immediately after natural defecation, ∼1–3 g of feces was scooped into sterile containers, frozen promptly (−201°C), and transferred to the laboratory for further storage (-80°C). All dogs were housed under standard domestic conditions (female dogs were kept both indoors and outdoors, while male dogs were housed exclusively outdoors. The dogs’ diet consisted of Gemon Lamb & Rice Adult (Monge, Italy), a nutritionally balanced dry food designed for adult dogs. Supplementary Table S3 contains the read counts and barcodes for the sequencing runs.

#### 3. Primer Comparison on Synthetic and Biological Samples

To evaluate three distinct 16S rRNA primer sets (ONT, MONT, and PB), we utilized:

1. Three Synthetic Mixtures:

`o` ZymoBIOMICS® Microbial Community Standard (Cat. #: D6300): This mock community comprises eight bacterial and two fungal strains, including both easy-to-lyse Gram-negative bacteria and tough-to-lyse Gram-positive bacteria and yeasts.
`o` ZymoBIOMICS® Gut Microbiome Standard (Cat. #: D6331): This standard features a more complex mock community comprised of 21 different strains, designed to mimic the human gut microbiome.
`o` ZymoBIOMICS® Fecal Reference with TruMatrix™ Technology (Cat. #: D6323): This reference material contains standardized human stool components, providing a complex microbial community representative of the human gut microbiome.

The reference composition used in our analyses is available under our GitHub repository https://github.com/Balays/CaniMeta/tree/main/standards

1. Two Biological Samples:

`o` One fecal sample from a 53-years old female volunteer (noninvasive, anonymous donation), sampled at Székesfehérvár, Hungary.
`o` One canine fecal sample (F5) collected under the same conditions as the others from Ajka-Padragkút, Hungary.

All mock standards were obtained from Zymo Research (Irvine, CA, USA). The biological fecal samples were placed on ice immediately after collection, then frozen at −201°C and later shipped to the lab for long-term storage (−801°C) and processing.

Each sample (synthetic or biological) was subjected to the three primer sets targeting the full 16S V1–V9 region, followed by Oxford Nanopore (MinION) or PacBio sequencing, as indicated.

### DNA purification

Depending on the study component, different DNA extraction protocols were applied as follows:

1) *Single-Canine Fecal Sample (Part*□*1):*

- Multiple isolation kits were tested, each using the manufacturer’s recommended input of 200 mg feces.
- For the Zymo High-Molecular-Weight (ZHMW) kit in particular, we adjusted to 100 mg of fecal material per the kit’s specific guidance for canine samples.
2) *Forty Dog Fecal Samples (Zymo HMW vs. MagBead, Part□2):*

- Each of the 40 targeted fecal samples was split into two aliquots. One aliquot underwent ZHMW extraction (using 100 mg of canine feces), while the other used the MagBead protocol at 200 mg, following the standard manufacturer’s recommendations.
3) *Mock Communities (MCS) and Additional Samples (Part*□*3):*

- In the evaluation of different 16S primer sets, 75 μL of the Zymo Microbial Community Standard (MCS) was used as input for each isolation kit, adhering to the manufacturer’s protocol.

All extractions were performed strictly according to each kit’s instructions, with the noted exceptions (100 mg vs. 200 mg feces for Zymo kits).

***Invitrogen™ PureLink™ Microbiome DNA Purification Kit*** The sample was mixed with 600 μl of S1 Lysis Buffer in a Bead Tube (provided in the kit) and homogenized by vortexing. Then, 100 µL of S2 Lysis Enhancer was added and the sample was vortexed again. The mix was incubated at 65°C for 10 minutes, followed by bead beating for homogenization using a vortex mixer set to maximum speed with horizontal agitation for 10 minutes. Afterward, the sample was centrifuged at 14,000 × g for 5 minutes. Subsequently, 400 µL of the clear supernatant was transferred to a new Eppendorf tube and combined with 250 µL of S3 Cleanup Buffer. The samples were centrifuged again at 14,000 × g for 2 minutes. Then, 500 µL of this supernatant was transferred to a clean tube and mixed with 900 µL of S4 Binding Buffer and briefly vortexed. Next, 700 µL of this mixture was loaded onto a spin column and centrifuged at 14,000 × g for 1 minute. The column was then placed into a new tube, and the remaining mixture was loaded for an additional 1-minute centrifugation at the same speed. The column was moved to a clean collection tube, and 500 µL of S5 Wash Buffer was added and centrifuged at 14,000 × g for 1 minute. To ensure all S5 Wash Buffer was removed, a final centrifugation was performed for 30 seconds at the same speed. Finally, the DNA from canine sample and from the MCS were eluted using 100 µL or 50 µL of S6 Elution Buffer from the spin columns, respectively. Samples were stored at-20°C.

***Macherey-Nagel™ NucleoSpin™ DNA Stool Mini Kit*** Samples were transferred to a Macherey-Nagel Bead Tube Type A, then, 850 µL of Buffer ST1 was added, and the samples were shaken horizontally for 3 seconds before being placed in a heat incubator. The samples were then incubated at 70°C for 5 minutes, followed by agitation on a Vortex-Genie® 2 at full speed and room temperature for 10 minutes, and then centrifuged at 13,000 x g for 3 minutes. Six hundred µL of the supernatant was transferred to a new 2 mL tube, and 100 µL of Buffer ST2 was added and briefly vortexed. The samples were incubated at 4°C for 5 minutes and centrifuged again at 13,000 x g for 3 minutes. Five hundred fifty µL of the lysate was then applied to a NucleoSpin® Inhibitor Removal Column and centrifuged for 1 minute at 13,000 x g, after which the column was discarded. Two hundred µL of Buffer ST3 was added to the samples and mixed thoroughly. Seven hundred µL of the mixture was then loaded onto a NucleoSpin® DNA Stool Column and centrifuged for 1 minute at 13,000 x g. The column was placed into a new tube for the washing process, which was repeated four times. Initially, 600 µL of Buffer ST3 was added to the column and centrifuged for 1 minute at 13,000 x g. The column was moved to a new tube, and 550 µL of Buffer ST4 was added and centrifuged for another minute at 13,000 x g. The column was transferred to a fresh tube, and 700 µL of Buffer ST5 was added, briefly vortexed, and then centrifuged for 1 minute at 13,000 x g. This step was repeated once more with 700 µL of Buffer ST5, followed by discarding the flow-through. The column was then dried by centrifugation for 2 minutes at 13,000 x g. Finally, 100 (for the dog sample) or 50 µL (for the MCS) of Buffer SE was pipetted directly onto the silica membrane of the column, and DNA was eluted by centrifuging for 1 minute at 13,000 x g. The DNA samples were subsequently stored at-20°C for future use.

***QIAGEN QIAamp Fast DNA Stool Mini Kit*** The stool sample (or the MCS mix) was placed in a 2 ml microcentrifuge tube and kept on ice. InhibitEX Buffer was added, and the sample was vortexed until fully homogenized. Large stool particles were removed via centrifugation. Next, 600 μl of the supernatant was combined with 25 μl of proteinase K and 600 μl of Buffer AL. After vortexing thoroughly, the mixture was heated to 95°C for 5 minutes (deviating from the kit’s recommended 70°C). Subsequently, 600 μl of 100% ethanol was added and mixed well. This lysate was then loaded onto a QIAamp spin column and centrifuged at 20,000 x g for 1 minute. The spin column was transferred to a new 2 ml tube, and the remaining lysate was loaded and centrifuged again. Following this, 500 μl of Buffer AW1 was added to the column and centrifuged at 20,000 x g for 1 minute before discarding the collection tube. After adding 500 μl of Buffer AW2 to the column and placing it in a new collection tube, the column was centrifuged at full speed for 3 minutes. To ensure no carryover of Buffer AW2, the column was then placed in a fresh 2 ml collection tube and centrifuged at full speed for an additional 3 minutes. The spin columns were transferred to clean Eppendorf tubes, and 100 µL of Buffer ATE was directly applied to the QIAamp membrane containing DNA from the canine sample. For the DNA derived from the MCS, we eluted it in 50 µL of Buffer ATE. After incubating at room temperature for 1 minute, the DNA was eluted with a final centrifugation at 20,000 x g for 1 minute into 50 μl of elution buffer and stored at-20°C.

***Zymo Research Quick-DNA™ HMW MagBead Kit*** The fecal sample was resuspended in 200 µl of DNA/RNA Shield™, followed by a 5-minute incubation at room temperature (20-30°C) on a tube rotator. Then, 33 µl of MagBinding Beads were added to each sample, mixed thoroughly, and shaken for 10 minutes. The sample was placed on a magnetic stand to allow the beads to separate clearly from the solution, and the supernatant was then removed. For the washing process, 500 µl of Quick-DNA™ MagBinding Buffer was added, the beads were resuspended, and the mixture was shaken for 5 minutes. The sample was placed back on the magnetic stand to discard the supernatant. Subsequently, 500 µl of DNA Pre-Wash Buffer was added and the beads were again resuspended. After returning the sample to the magnetic stand and discarding the supernatant, 900 µl of g-DNA Wash Buffer was added, mixed, and transferred to a new tube. The magnetic stand facilitated the separation of the beads from the solution, and the supernatant was discarded. This washing step was repeated once more. The beads were allowed to air dry for 20 minutes. In the elution phase, 50 µl of DNA Elution Buffer was added to the beads. Following mixing, the solution was left to incubate at room temperature for 5 minutes. The sample was then placed on the magnetic stand once more to separate the beads from the solution. The eluted DNA was carefully transferred to a new tube and stored at-20°C for later analysis.

***Zymo Research ZymoBIOMICS™ 96 MagBead DNA Kit - Dx*** Samples were added to the Zymo Research (ZR) BashingBead™ Lysis Tubes and 750 µl of ZymoBIOMICS™ Lysis Solution was also added. The mixture was then centrifuged at ≥ 10,000 x g for 1 minute. Two hundred µl of the supernatant were transferred to new tubes, and then 600 µl of ZymoBIOMICS™ MagBinding Buffer was added. Twenty-five µl of ZymoBIOMICS™ MagBinding Beads were dispensed into each tube, and the contents were mixed on a shaker plate for 10 minutes. The tubes were then placed on a magnetic stand until the beads settled, after which the supernatant was aspirated and discarded. Tubes were removed from the magnet, and 500 µl of ZymoBIOMICS™ MagBinding Buffer was added and mixed well on a shaker plate for 1 minute. The process of placing the tubes on the magnetic stand to settle the beads and discarding the supernatant was repeated. Subsequently, 500 µl of ZymoBIOMICS™ MagWash 1 was added to each tube and shaken for 1 minute. After placing the tubes on the magnet again and discarding the supernatant, 900 µl of ZymoBIOMICS™ MagWash 2 was added and mixed thoroughly for 1 minute. The tubes were placed on the magnet until the beads settled and the supernatant was again discarded. The final washing step was repeated. The samples were heated on a thermal block at 55°C until the beads dried, which took approximately 10 minutes. Then, 50 µl of ZymoBIOMICS™ DNase/RNase-Free Water was added to each tube and the beads were resuspended. The mixture was shaken for 10 minutes, then transferred to a magnetic stand for 2-3 minutes until the beads settled. The supernatant containing the eluted DNA was transferred to a clean elution tube. The eluted DNA samples were stored at-20°C for future use.

### Library preparation

Depending on the study objective—whether metagenomic whole-genome sequencing (mWGS) or targeted 16S rRNA amplicon sequencing—different library preparation kits and sequencing platforms were employed:

1) *Metagenomic Whole-Genome Sequencing (mWGS)*

- **Illumina Platforms (MiSeq and NovaSeq)**: Libraries were constructed using the Illumina DNA Prep Kit for the MiSeq and the xGen™ DNA Library Prep Kit (IDT) for the NovaSeq runs. These short-read libraries targeted ∼1300–5001bp inserts and underwent paired-end sequencing.
- **Oxford Nanopore Technologies (ONT, MinION)**: For long-read mWGS, we used the ONT Native Barcoding Kit 24 V14, enabling multiplexed sequencing of canine fecal DNA on the MinION device.
2) *16S rRNA Gene Amplicon Sequencing (Full V1–V9 Region)*

- **PacBio Sequellle**: Amplicon libraries covering the V1–V9 region were prepared following the PacBio full-length 16S protocol (e.g., barcoded adapters for multiplexing).
- **Oxford Nanopore (MinION)**: Similarly, V1–V9 libraries were constructed with ONT’s 16S barcoding kit (standard or modified primer sets), allowing comparisons to the PacBio results.
3) *Partial 16S Amplicon Sequencing (Selected V Regions on Illumina)*

- For subsets of samples, V1–V2, V3–V4, or V1–V3 amplicons were generated.
- Standard Illumina 16S workflows (e.g., Nextera-style adapters) were used, producing paired-end reads (∼300 bp) on the MiSeq.

All library preparations followed the manufacturers’ protocols, with routine QC checks (Qubit quantification, TapeStation or Bioanalyzer) confirming DNA concentration and fragment size distributions.

#### IDT xGen™ DNA Lib Prep Kit

DNA samples were adjusted to a final volume of 19.5 µL using Low EDTA TE buffer. Components including 3 µL of Buffer K1, 1.5 µL of Reagent K2, and 6 µL of Enzyme K3 were added to each sample. The mixture was then subjected to enzymatic processing in a thermal cycler programmed with the Enzymatic Prep settings: fragmentation at 32°C for a specific duration (time not specified), followed by inactivation at 65°C for 30 minutes, with the thermal cycler lid heated to 70°C. Post-reaction, samples were stored at 4°C. The fragmented DNA underwent ligation preparation by adding 12 µL of Buffer W1, 4 µL of Enzyme W3, 5 µL of stubby adapter Reagent W5, and 9 µL of Low EDTA TE to each sample, making up a 30 µL Ligation Master Mix. Ligation was conducted at 20°C for 20 minutes. Post-ligation, the samples were cleaned up using SPRIselect beads. Initially, 48 µL of beads were added to each sample at room temperature, mixed by vortexing for 5 seconds, and briefly centrifuged. The samples were incubated for 5 minutes at room temperature and then placed on a magnetic rack until the supernatant cleared. The supernatant was discarded, and the beads were washed twice with 180 µL of freshly prepared 80% ethanol, each wash followed by a brief incubation and removal of ethanol. After drying, 20 µL of Low EDTA TE was added, and the beads were resuspended. DNA was eluted into a new tube after a 2-minute incubation at room temperature. PCR amplification was then performed on the eluted DNA. The PCR setup involved an initial denaturation at 98°C for 45 seconds, followed by cycles of denaturation at 98°C for 15 seconds, annealing at 60°C for 30 seconds, and extension at 72°C for 30 seconds, concluding with a final extension at 72°C for 1 minute. A final post-PCR cleanup was conducted similar to the initial cleanup steps, using 48 µL of beads per sample and repeating the washing steps with 80% ethanol. The final DNA was resuspended in 21 µL of Low EDTA TE and transferred to a clean tube for storage or further analysis.

#### Illumina DNA Prep

Sixty nanograms of DNA (in a 30 µl volume) were used for each sample. A mix of 10 µl Tagmentation Buffer 1 (TB1) and 10 µl Bead-Linked Transposomes (BLT) was prepared, and 20 µl of this mix was added to the DNA. The sample was then incubated at 55°C for 15 minutes before being cooled to 10°C. Subsequently, 10 µl of Tagment Stop Buffer (TSB) was added to the mixture and gently combined. The sample was incubated at 37°C for 15 minutes, followed by cooling to 10°C. The samples were then placed on a magnetic stand for 3 minutes to separate the beads, and the supernatant was discarded. The beads were then washed by slowly adding 100 µl of Tagment Wash Buffer (TWB) after removing the sample from the magnet. The sample was returned to the magnetic stand, the wash was repeated, and the supernatant discarded again. A mix consisting of 20 µl Enhanced PCR Mix (EPM) and 20 µl of nuclease-free water was prepared, and 40 µl of this mix was added to the beads. Index adapters (i5 and i7, 5 µl each; Table 2) were added, and PCR was performed as follows: the reaction was pre-heated at 68°C for 3 minutes, followed by an initial denaturation at 98°C for 3 minutes. This was succeeded by 5 cycles of denaturation at 98°C for 45 seconds, annealing at 62°C for 30 seconds, and elongation at 68°C for 2 minutes. A final extension was then performed at 68°C for 1 minute. Post-amplification, the libraries underwent a cleanup process. Initially, the samples were placed on a magnetic stand for about 5 minutes. Forty-five µl of the PCR product’s supernatant was transferred into a new tube. To this supernatant, 40 µl of nuclease-free water and 45 µl of Sample Purification Beads (SPB) were added and mixed at 1600 rpm for 1 minute, followed by a 5-minute room temperature incubation. The samples were then placed on the magnet, and 125 µl of the supernatant was transferred to new tubes containing 15 µl of undiluted SPB. The samples were mixed at 1600 rpm for 1 minute and incubated at room temperature for another 5 minutes. Post-mixing, the supernatant was discarded, and the samples were washed twice with 200 µl of 80% ethanol, each time incubating the sample on the magnetic stand for 30 seconds to allow ethanol removal. After the final wash, the pellet was air-dried, and then 32 µl of RSB was added to resuspend the beads. This mixture was incubated for 2 minutes, placed on the magnetic stand for another 2 minutes, and finally, the supernatant containing the prepared library was transferred to a new tube.

#### Oxford Nanopore Technologies Ligation Sequencing DNA V14 Kit

One μg of genomic DNA was adjusted the volume up to 48 μl and added 3,5 μl NEBNext FFPE DNA Repair Buffer, 3,5 μl Ultra II End-prep Reaction Buffer, 3 μl Ultra II End-prep Enzyme Mix, and 2 μl NEBNext FFPE DNA Repair Mix, all from New England Biolabs (NEB). The mixture was incubated in a thermal cycler at 20°C for 5 minutes then 65°C for 5 minutes. After the incubation, AMPure XP beads was resuspended and 60 μl of the bead suspension was added to the end-prep reaction. The sample was incubated for 5 minutes on a Hula mixer at room temperature then washed with 80% ethanol. After the ending-repair, the DNA was eluted in 61 μl of nuclease-free water. For adapter ligation, 5 μl Ligation Adapter, 10 μl NEBNext Quick T4 DNA Ligase, and 25 μl Ligation Buffer was added to the sample. After 10 minutes of incubation at RT, 40 μl of resuspended AMPure XP Beads were added to the reaction. The mixture was incubated on a Hula mixer for 10 minutes, then washed with Short Fragment Buffer (SFB), and the DNA was eluted in 13 μl of Elution Buffer.

#### Combining the ONT Rapid 16S Barcoding Kit and Native Barcoding Kit for Custom Primer Sequences

Because the Oxford Nanopore Technologies Rapid Sequencing 16S Barcoding Kit incorporates barcoded primers, a combined protocol was employed to enable the attachment of sequencing barcodes and adapters to the custom or modified primer sequences.

According to the Rapid Sequencing Amplicons protocol (SQK-16S024), for each sample 10 ng genomic DNA was used in 10µL of nuclease-free water. 5 µL nuclease-free water, 10 µL input DNA (10 ng), and 25 µL of the PCR master mix (LongAmp Hot Start Taq 2× Master Mix, New England BioLabs, USA) were mixed in a 0.2 mL thin-walled PCR tubes. We used three different primer sets which were the ONT primer (forward: 5’ - AGAGTTTGATCMTGGCTCAG - 3’, reverse: 5’ - CGGTTACCTTGTTACGACTT - 3’), modified ONT primer (forward: 5’ - AGRGTTYGATYMTGGCTCAG-31, reverse: 5’ - CGGYTACCTTGTTACGACTT-31) and the PacBIO 16S rRNA degenerate primer (forward: 5’ - AGRGTTYGATYMTGGCTCAG - 3’, reverse: 5’ - RGYTACCTTGTTACGACTT - 3’). 10 µL of each primer pair was added to each sample replicate (5 µL forward + 5 µL reverse) and mixed thoroughly. In a thermal cycler, the reaction was amplified using the following cycling conditions: initial denaturation 1 min (95°C, 1 cycle), denaturation 20 secs (95°C, 25 cycles), annealing 30 secs (55°C, 25 cycles), extension 2 mins (65°C, 25 cycles), final extension 5 mins (65°C, 1 cycle), and hold on 4°C. On a magnetic rack, the PCR products were cleaned using AMPure XP beads, washed with 70% ethanol, and eluted in 10 µL nuclease-free water. From the following step, we used the Native Barcoding Kit.

We used 200 fmol amplicon DNA and made up each sample to 12,5 μl with Nuclease-free water than added Ultra II End-prep Reaction Buffer and Ultra II End-prep Enzyme Mix from New England Biolabs (NEB). The mixture was well mixed, briefly centrifuged, and then incubated in a thermal cycler at 20°C for 5 minutes followed by 65°C for 5 minutes. 0.75 μl of samples were transferred to a new strip. The NEB Blunt/TA Ligase Master Mix was prepared as per the manufacturer’s instructions and kept on ice. After thawing and mixing, the reagents were spun do Native Barcodes (NB01-45) were prepared and kept on ice. To each sample, a selected Native Barcode, Blunt/TA Ligase Master Mix and Nuclease-free water were added. The mixture was then incubated for 20 minutes at room temperature. After that EDTA was added to stop the reaction and the barcoded samples were pooled.

The pooled sample was mixed with 0.4X volume of AMPure XP Beads and incubated on a Hula mixer for 10 minutes. After washing with 80% ethanol, DNA was eluted in 31 μl of nuclease-free water, with intermittent flicking at 37°C to facilitate elution.

For adapter ligation, the NEBNext Quick Ligation Reaction Module was prepared according to the manufacturer’s instructions. The pooled barcoded sample was combined with Native Adapter, Quick T4 DNA Ligase, and NEBNext Quick Ligation Reaction Buffer (5X). Following ligation, 20 μl of AMPure

XP Beads were added to the reaction. The mixture was incubated on a Hula mixer for 10 minutes, washed with Short Fragment Buffer (SFB), and the DNA was eluted in 15 μl of Elution Buffer after a 10-minute incubation at 37°C. The final DNA library was prepared by adjusting the concentration to the desired level with Elution Buffer for sequencing.

#### Oxford Nanopore Technologies Rapid Sequencing 16S Barcoding Kit

Ten ng of high molecular weight genomic DNA, in a 10 µl volume, was utilized for library preparation from both the canine and MCS samples. However, DNA isolated using the QIAGEN kit did not meet this high molecular weight criterion. The input DNA was combined with 14 µl of nuclease-free water (Invitrogen), 1 µl of 16S Barcode (1 µM; Table 3, Table 4), and 25 µl of LongAmp Taq 2X master mix (New England Biolabs). The PCR amplification of the samples was performed starting with an initial denaturation at 95°C for 1 minute. This was followed by 25 cycles of denaturation at 95°C for 20 seconds, annealing at 55°C for 30 seconds, and extension at 65°C for 2 minutes. A final extension was carried out at 65°C for 5 minutes to complete the reaction. Post-amplification, the DNA samples were transferred to clean 1.5 ml Eppendorf DNA LoBind tubes and mixed with 30 µl of resuspended AMPure XP beads. The samples were then agitated on a Hula mixer for 5 minutes at room temperature. Afterward, the tubes were placed on a magnetic rack, and the supernatant was discarded. The beads were washed with 200 µl of freshly prepared 70% ethanol, the ethanol was removed, and the washing step was repeated. Following air drying of the beads, the tubes were removed from the magnet, and the beads were resuspended in 10 µl of 10 mM Tris-HCl pH 8.0 with 50 mM NaCl. After a 2-minute incubation at room temperature, the tubes were placed back on the magnet. Ten µl of the clean supernatant, containing the ONT libraries, was transferred to a new Eppendorf DNA LoBind tube. The barcoded libraries were then pooled in an equal molar ratio, and 1 µl of RAP was added. The mixture was incubated for 5 minutes at room temperature. Subsequently, one hundred femtomoles of the libraries were loaded onto a MinION flow cell for sequencing.

#### Pacific Biosciences Full-Length 16S Library Preparation Using SMRTbell Express Template Prep Kit

***2.0*** For each sample, a mixture was prepared consisting of 1.5 µL of PCR-grade water and 12.5 µL of 2X KAPA HiFi HotStart ReadyMix. To this mixture, 3 µL of a barcoded forward primer solution (2.5 µM, primer identification numbers are detailed in Table 5) was added, followed by 3 µL of the corresponding reverse primer (also listed in Table 5) and 5 µL of the DNA sample. The DNA was then amplified according to the following protocol: initial denaturation at 95°C for 3 minutes, followed by 25 cycles of denaturation at 95°C for 30 seconds, annealing at 57°C for 30 seconds, and extension at 72°C for 60 seconds.

#### PerkinElmer NEXTFLEX® 16S V1-V3 Amplicon-Seq Kit for Illumina

Genomic DNA, with concentrations ranging from 1.6 ng to 36 ng as noted in Table 6, was diluted to a total volume of no more than 36 µL using nuclease-free water. Then, 12 µL of NEXTflex™ PCR Master Mix and 2 µL of the 16S V1-V3 PCR I Primer Mix were added to the solution. The final reaction volume was adjusted to 50 µL. The first step of the PCR amplification was performed as follows: an initial denaturation at 98°C for 4 minutes, followed by 8 cycles of 30 seconds denaturation at 98°C, 30 seconds annealing at 60°C, and 30 seconds extension at 72°C. The process concluded with a final extension at 72°C for 4 minutes. PCR cleanup involved adding 50 µL of AMPure XP Beads (Beckman Coulter) to each sample. After thorough mixing, the samples were left to incubate at room temperature for 5 minutes. They were then placed on a magnetic stand until the supernatant became clear. The supernatant was discarded, and the beads were washed twice with 200 µL of freshly prepared 80% ethanol. The samples were air-dried for 3 minutes and then resuspended in 38 µL of Resuspension Buffer. After incubating for an additional 2 minutes at room temperature, 36 µL of the clear supernatant was transferred to fresh tubes. Additional PCR amplification was conducted by adding 12 µL of NEXTflex™ PCR Master Mix and 2 µL of NEXTflex™ PCR II Barcoded Primer Mix, under the following conditions: an initial denaturation at 98°C for 4 minutes, followed by a variable number of cycles depending on the initial DNA quantity. Each cycle consisted of 30 seconds at 98°C for denaturation, 30 seconds at 60°C for annealing, and 30 seconds at 72°C for extension. The reaction was completed with a final extension step at 72°C for 4 minutes. The PCR cleanup was completed as per the guidelines provided for post-first PCR purification.

#### Zymo Research Quick-16S™ NGS Library Prep Kit

Ten µl of Quick-16S™ qPCR Premix were combined with 4 µl of Quick-16S™ Primer Set V1-V2 or V3-V4 and 4 µl of ZymoBIOMICS® DNase/RNase Free Water. Additionally, 2 µl of DNA samples (2.5 ng/µl) were incorporated. PCR was executed in a Verity Thermal Cycler (Applied Biosystems) following the guidelines provided in the Zymo Research Manual as follows: initial denaturation at 95°C for 10 minutes, followed by 20 cycles for canine samples and 12 cycles for MCS samples with denaturation at 95°C for 30 seconds, annealing at 55°C for 30 seconds, and extension at 72°C for 3 minutes. After amplification, 1 µl of Reaction Clean-up Solution was added to each sample, which was then incubated at 37°C for 15 minutes. The reactions were halted by heating to 95°C for 10 minutes, after which the samples were cooled to 4°C. Subsequently, 10 µl of Quick-16S™ qPCR Premix and 4 µl of ZymoBIOMICS® DNase/RNase Free Water were mixed. Index primers (2 µl each from ZA5 and ZA7, with detailed pairs and sequences found in Table 7 and 8) along with 2 µl of the amplified DNA were added to the mixture. Barcoded PCR reactions were conducted according to the manual’s recommendations: an initial denaturation at 95°C for 10 minutes, followed by 5 cycles consisting of 30 seconds at 95°C for denaturation, 30 seconds at 55°C for annealing, and a 3-minute extension at 72°C. For the purification of the PCR products, Select-a-Size MagBeads were utilized. Initially, the MagBeads were agitated to resuspend, and 16 µl were mixed with each sample. This mixture was then allowed to sit at room temperature for 5 minutes before being placed on a magnetic rack for 3 to 10 minutes. The supernatant was discarded, and the beads were washed twice with 200 µl of DNA Wash Buffer. The samples were briefly removed from the magnetic field and allowed to sit for 3 minutes at room temperature to ensure complete buffer removal. Libraries were then eluted in 25 µl of DNA Elution Buffer and stored at-20°C for subsequent analysis.

#### Quality and Quantity Check

After the DNA isolation and library preparation methods (Oxford Nanopore, Illumina MiSeq), the genomic DNA samples were quantified and assessed for quality. DNA concentrations were measured using the Invitrogen™ Qubit™ 4 Fluorometer with the Qubit™ 1X dsDNA High Sensitivity (HS) kit. For quality assessment, the Agilent Technologies 4150 TapeStation System was used. The Genomic DNA ScreenTape was applied for genomic DNA and Oxford Nanopore WGS evaluation, the D5000 ScreenTape for ONT V1-V9, and the D1000 ScreenTape for V region amplicon libraries.

## Bioinformatics

### Single-Canine, Multi-Platform Dataset

#### Pre-processing of NovaSeq Data

The raw reads were pre-processed using fastp [19], using the following parameters: *--trim_front1 12--trim_front2 12 trim_tail1 3 trim_tail2 3-q 25-p-P 100--dedup--merge--correction--trim_poly_g.* This involved trimming adapters, merging overlapping reads, filtering by quality, deduplicating, correcting errors, and trimming polyG sequences. The raw.fastq files – R1 and R2 – were submitted to the European Nucleotide Archive (ENA) under the following Project ID: **PRJEB75753,** and accessions: **ERR13110291** and **ERR13110292**, respectively.

#### Pre-processing of MinION Data

Raw sequencing data from the Oxford Nanopore Technologies (ONT) MinION platform were basecalled using dorado [20] with the’super accurate’ model, utilizing an R9 flow cell. The resulting.fastq file were submitted to ENA and be accessed using the following link: Accession: **ERR14242610, Project ID: PRJEB75753**

#### Taxonomic Classification

For both our new sequencing dataset (ONT/NovaSeq) and the previously published MiSeq reads (available under **PRJEB59610**), taxonomic classification was first performed using sourmash v4 [21], utilizing its combined database spanning bacteria, archaea, eukaryotes, and viruses. Sourmash utilizes FracMinHash sketching to efficiently represent and compare large genomic and metagenomic datasets by compressing sequences into smaller, representative sketches, enabling rapid similarity searches and analyses. It was shown in [17] that it provides good accuracy and precision for both long and short read metagenomic sequencing datasets.

Subsequently, to focus on viral composition, we utilized Kraken 2 [22] with a dedicated viral database. Kraken 2 assigns taxonomic labels by examining k-mers within each query sequence; its probabilistic hash-table structure offers faster classification speeds and lower memory usage compared to earlier methods. After classification, **kreport2krona.py** from **KrakenTools** [23] was employed to convert Kraken 2 outputs into abundance files for visualization and downstream analyses.

### Pre-processing and Taxonomic Classification for the DNA Method Comparison on Forty Canine Samples Dataset

Raw sequencing data from the Oxford Nanopore Technologies (ONT) MinION platform were basecalled using Dorado (version 0.7.3.9) with the’super accurate’ model, utilizing an R9 flow cell. Following basecalling, reads were demultiplexed based on their barcodes. The resulting.fastq file were submitted to the ENA and be accessed using the following link: **PRJEB82097.**

For taxonomic classification, full-length 16S rRNA reads were analyzed using Emu with standard parameters. Emu employs an expectation-maximization algorithm to generate taxonomic abundance profiles from full-length 16S rRNA reads, effectively handling the higher error rates associated with long-read sequencing technologies [24].

### Pre-processing and Taxonomic Classification of the V1-V9 Primer Comparison on Diverse Samples Dataset

The same bioinformatic workflow was utilized as in the *Forty Dog Fecal Samples* section, with the exception that this sequencing was carried out on the R10 flow-cell. The.fastq file were submitted to ENA and **for** the synthetic and canine samples can be accessed using the following link: **PRJEB85420,** and for the human samples**: PRJEB85714**.

### Downstream Data Processing

#### Data Import

The output of each bioinformatic method was converted into phyloseq-type R objects [25], containing sample information (metadata) and taxonomic composition (and lineage). Subsequently, all downstream data processing was carried out within the R-enviroment, utilizing our in-house developed collection of R scripts.

#### Correlation between ZHMW and ZMB kits in the Forty Dog Fecal Samples Dataset

To assess the consistency between DNA isolation methods, we computed correlations (R² values) between microbial compositions obtained from the ZHMW and ZMB kits. A linear regression model was fitted for each sample and each taxon level and the correlations were visualized.

#### Statistics, Beta diversity

Beta diversity analyses were performed to investigate differences in microbial community composition among samples. Non-metric multidimensional scaling (NMDS) ordination was conducted using Bray–Curtis dissimilarity, and the phyloseq package [25] in R was employed to generate distance matrices. In addition, principal coordinates analysis (PCoA) was carried out on the same Bray–Curtis distance matrix, providing an alternative ordination view of sample clustering.

To assess the effects of DNA isolation method, sex, and sampling date on community structure, we used permutational multivariate analysis of variance (PERMANOVA; adonis2, vegan package [26]). For the Forty Dog Fecal Samples Dataset, we accounted for repeated measures by specifying the dog ID as a stratification factor (strata) within adonis2. In the Single Canine Fecal Sample Dataset, PERMANOVA modeled microbial composition as a function of both DNA isolation method and library preparation approach, again accounting for repeated treatments of the same fecal sample.

Additionally, the homogeneity of dispersion among groups was tested using PERMDISP2 (betadisper, vegan). This analysis determines whether variance within each group significantly differs, potentially affecting beta diversity comparisons. The significance of these dispersion differences was evaluated with a permutation test (permutest), and Tukey’s honest significant difference (TukeyHSD) was used post hoc to identify pairwise group differences. Ordination plots (PCoA axes or NMDS dimensions) were visualized using ggplot2 [27], with centroids and confidence ellipses (95% level) drawn to illustrate how tightly (or loosely) samples within each group cluster. Boxplots of distances to centroid were also generated to highlight group-specific dispersion results.

#### Visualization

Data were visualized using mainly using bar plots for taxonomic composition summaries, PERMANOVA term contribution summaries, and line plots for correlation values, utilizing ggplot2 [27].

## Abbreviations

BLT: Bead-Linked Transposomes
EPM: Enhanced PCR Mix
ENA: European Nucleotide Archive,
FR: fecal reference
HMW: high molecular weight
I: Invitrogen PureLink™ Microbiome DNA Purification Kit
LRS: long-read sequencing
M: Macherey-Nagel NucleoSpin® DNA Stool Mini Kit
MCS: Mock Communities
MONT: modified ONT
mWGS: metagenomic whole-genome sequencing
NEB: New England Biolabs
NMDS: Non-metric multidimensional scaling
ONT: Oxford Nanopore Technologies
PERMANOVA: Permutational multivariate analysis of variance
PERMDISP: Permutational multivariate analysis of dispersion
PB: PacBio
PCoA: Principal Coordinate Analysis
Q: Qiagen QIAamp Fast DNA Stool Mini Kit
SFB: Short Fragment Buffer
SRS: short-read sequencing
SPB: Sample Purification Beads
TB1: Tagmentation Buffer 1
TSB: Tagment Stop Buffer
TWB: Tagment Wash Buffer
WGS: whole-genome shotgun
Z: Zymo Research Quick-DNA™ HMW MagBead Kit
ZHMW: Zymo High-Molecular-Weight
ZMB: Zymo MagBead
ZR: the Zymo Research

## Data Availability

### Single-Canine, Multi-Platform Dataset

All raw FASTQ reads and associated metadata have been submitted to the ENA under the accession **PRJEB59610** and **PRJEB75753** for the.

### DNA Method Comparison on Forty Canine Samples

All raw FASTQ reads and associated metadata have been submitted to the ENA under the accession **PRJEB82097**.

### V1-V9 Primer Comparison on Diverse Samples

All raw FASTQ reads and associated metadata have been submitted to the ENA under the accession: **PRJEB85420** for the synthetic and canine samples and **PRJEB85714** for the human samples.

## Code Availability

All scripts used for data analysis and visualization are publicly available in the GitHub repository https://github.com/Balays/CaniMeta. The R Markdown files included in the repository allow for full reproducibility of the workflow, including all analyses and visualizations.

## Ethical Approval

This study was conducted in accordance with the principles of the Declaration of Helsinki. Ethical approval was obtained from the Medical Research Council, Budapest, Hungary, under the accession number BMEÜ/725-1 /2022/EKU. Written informed consent was obtained from the participant for sample collection and data publication.

## Acknowledgements

We thank the Tombácz family from Szeged, Hungary, the owners of Toto, for their support in sample collection. We express our gratitude to Gabriella Kassai, owner of the Serteperti Pumi Kennel [Kiskunmajsa (formerly Dány), Hungary; https://serteperti.hu/], for providing samples from their dogs (Degesz, Ilka, Sampo, Lonci, Ajsa, Matyi, Rozi, and Mini), and to Melinda Takácsné Horváth, the owner of the Rezeta-Réti Pumi Kennel (Ajka-Padragkút, Hungary; https://www.facebook.com/p/Rezerta-R%C3%A9ti-PUMI-Kennel-100057382122448/), for providing Hanga’s sample.

## Author contributions

D.T. and Z.B. conceived and designed the experiments. Á.D., A.T. and T.J. isolated the DNA. Á.D., A.T., and T.J. prepared the sequencing libraries and performed sequencing. B.K., G.G., D.T., and Z.B. analyzed the data. I.P., T.J., Á.D. and G.G. handled the dataset. B.K., Z.B. and D.T. wrote the manuscript. D.T. supervised the project. All authors read and approved the final version of the manuscript.

## Legends for Supplementary Figures and Tables

**Supplementary Figure S1.** Bar plots of *R*˄2 values from PERMANOVA analyses evaluating how “DNA Isolation Method” and “Library Preparation Protocol” influence variation in the Dog_M0 V-region dataset. **(A)** Results from the full analysis, including all DNA extraction kits and library protocols. **(B)** An identical analysis excluding Qiagen-extracted samples. Each bar indicates the proportion of community dissimilarity (*distance*) attributed to a particular factor, with numeric labels showing the exact *R*˄2 values. The residual bar captures unaccounted variation in the model.

**Supplementary Figure S2.** Stacked bar plots showing the top 30 virus composition across Dog_M0 WGS runs (including MySeq, ONT, and NovaSeq data) classified using Kraken12’s viral database. The Y-axis represents the proportion of reads assigned to each viral species, with unclassified and non-viral reads excluded. Unfilled portion in each bar represents taxa outside the top130.

**Supplementary Table S1.** Overview of the Pumi dogs included in the study, detailing their age, sampling dates, reproductive status, and living conditions at the time of sampling.

**Supplementary Table S2.** Read counts and other information on the Illumina 16S and ONT 16S V-region sequencing data from the single-canine (Dog_M0) sample

**Supplementary Table S3.** Read counts and other information on the Illumina NovaSeq WGS and ONT WGS sequencing data from the single-canine (Dog_M0) sample

**Supplementary Table S4.** Read counts and other information on the ZMB1ZHMW DNA isolation kit comparison on the Serteperti dog cohort.

**Supplementary Table S5.** Read counts and other information on the primer comparison on the five different samples.

