## Supplementary figures and images for "Comprehensive Metagenomic Profiling of Diverse Microbiomes"

### Supplementary Figure S1

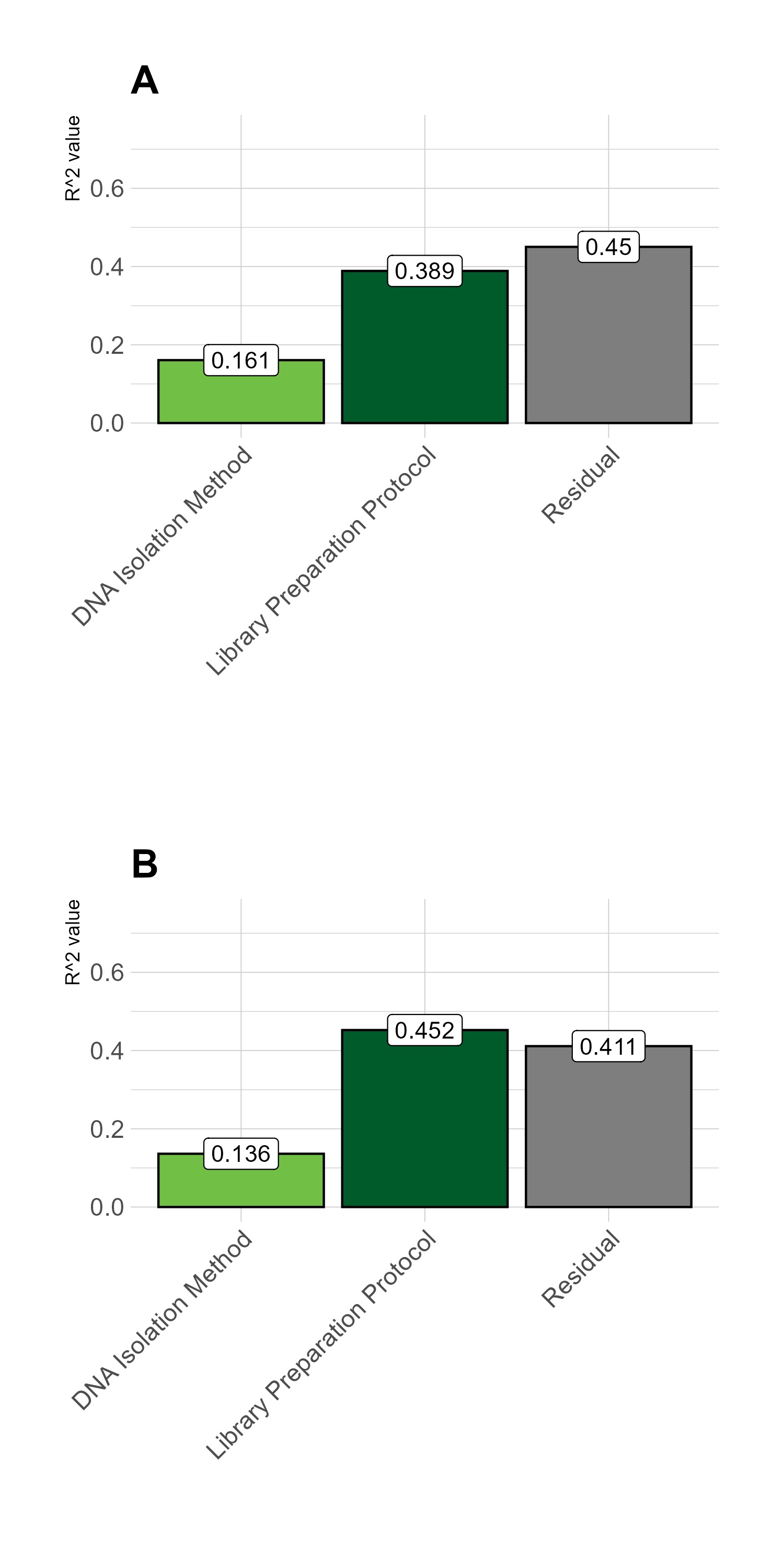

### Supplementary Figure S2

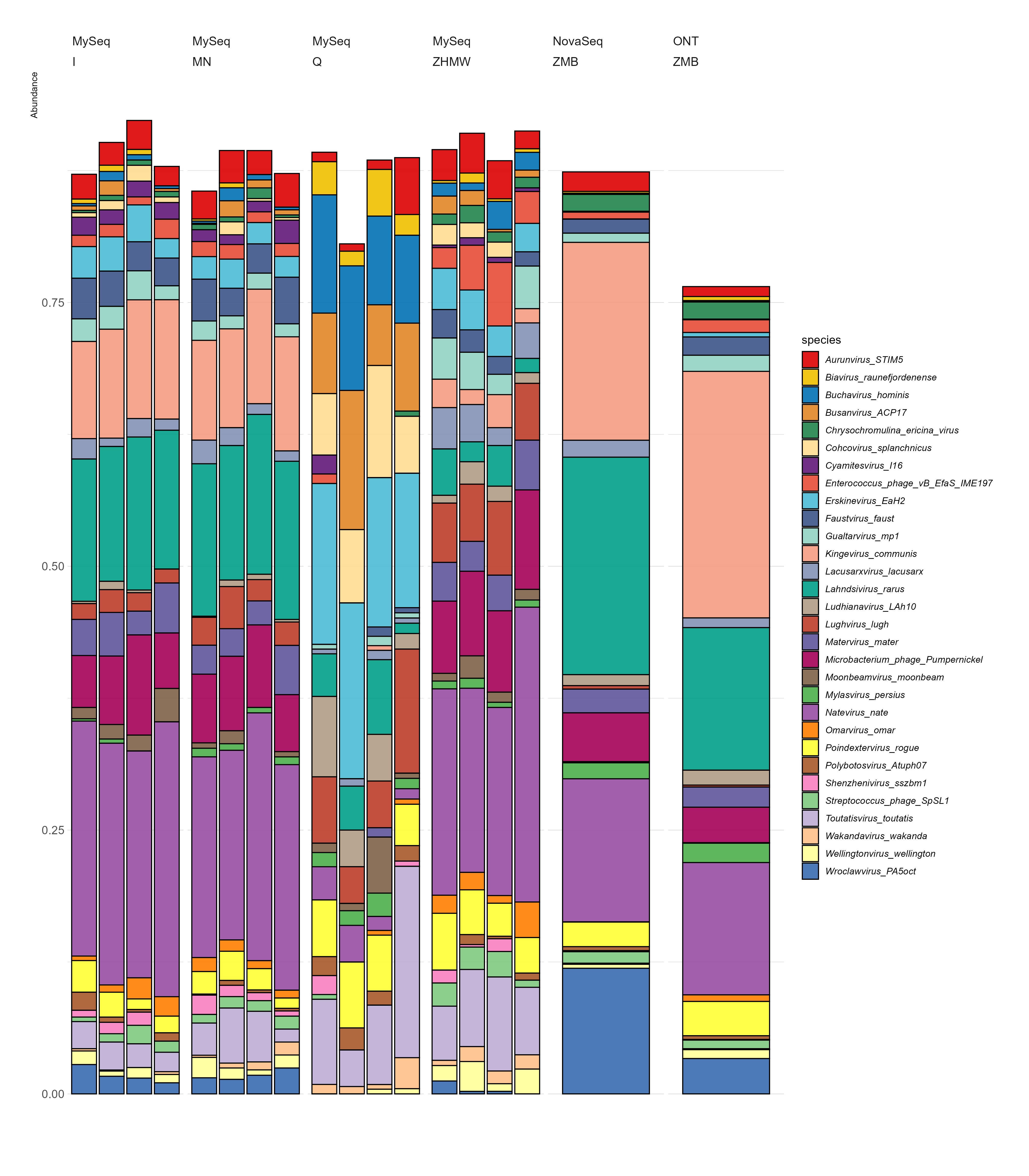
